# Consensus transcriptional regulatory networks of coronavirus-infected human cells

**DOI:** 10.1101/2020.04.24.059527

**Authors:** Scott A Ochsner, Rudolf T Pillich, Neil J McKenna

## Abstract

Establishing consensus around the transcriptional interface between coronavirus (CoV) infection and human cellular signaling pathways can catalyze the development of novel anti-CoV therapeutics. Here, we used publicly archived transcriptomic datasets to compute consensus regulatory signatures, or consensomes, that rank human genes based on their rates of differential expression in MERS-CoV (MERS), SARS-CoV-1 (SARS1) and SARS-CoV-2 (SARS2)-infected cells. Validating the CoV consensomes, we show that high confidence transcriptional targets (HCTs) of CoV infection intersect with HCTs of signaling pathway nodes with known roles in CoV infection. Among a series of novel use cases, we gather evidence for hypotheses that SARS2 infection efficiently represses E2F family target genes encoding key drivers of DNA replication and the cell cycle; that progesterone receptor signaling antagonizes SARS2-induced inflammatory signaling in the airway epithelium; and that SARS2 HCTs are enriched for genes involved in epithelial to mesenchymal transition. The CoV infection consensomes and HCT intersection analyses are freely accessible through the Signaling Pathways Project knowledgebase, and as Cytoscape-style networks in the Network Data Exchange repository.

## Introduction

Infection of humans by coronaviruses (CoV) represents a major current global public health concern. Signaling within and between airway epithelial and immune cells in response to infections by CoV and other viruses is coordinated by a complex network of signaling pathway nodes. These include chemokine and cytokine-activated receptors, signaling enzymes and transcription factors, and the genomic targets encoding their downstream effectors^1–3^. Placing the transcriptional events resulting from CoV infection in context with those associated with host signaling paradigms has the potential to catalyze the development of novel therapeutic approaches. The CoV research community has been active in generating and archiving transcriptomic datasets documenting the transcriptional response of human cells to infection by the three major CoV strains, namely, Middle East respiratory syndrome coronavirus (MERS-CoV, or MERS) and severe acute respiratory syndrome coronaviruses 1 (SARS-CoV-1, or SARS1) and 2 (SARS-CoV-2, or SARS2)^4–9^. To date however the field has lacked a resource that fully capitalizes on these datasets by, firstly, using them to identify human genes that are most consistently transcriptionally responsive to CoV infection and secondly, contextualizing these transcriptional responses by integrating them with ‘omics data points relevant to host cellular signaling pathways.

We recently described the Signaling Pathways Project (SPP)^10^, an integrated ‘omics knowledgebase designed to assist bench researchers in leveraging publically archived transcriptomic and ChIP-Seq datasets to generate research hypotheses. A unique aspect of SPP is its collection of consensus regulatory signatures, or consensomes, which rank genes based on the frequency of their significant differential expression across transcriptomic experiments mapped to a specific signaling pathway node or node family. By surveying across multiple independent datasets, we have shown that consensomes recapitulate pathway node-genomic target regulatory relationships to a high confidence level^10^. Here, as a service to the research community to catalyze the development of novel CoV therapeutics, we generated consensomes for infection of human cells by MERS, SARS1 and SARS2 CoVs. Computing the CoV consensomes against those for a broad range of cellular signaling pathway nodes, we discovered robust intersections between genes with high rankings in the CoV consensomes and those of nodes with known roles in the response to CoV infection. Integration of the CoV consensomes with the existing universes of SPP transcriptomic and ChIP-Seq data points in a series of use cases illuminates previously uncharacterized interfaces between CoV infection and human cellular signaling pathways. Moreover, while this paper was under review and revision, numerous contemporaneous and independent wet bench-based studies came to light that corroborate *in silico* predictions made using our analysis pipeline. To reach the broadest possible audience of experimentalists, the results of our analysis were made available in the SPP website, as well as in the Network Data Exchange (NDEx) repository. Collectively, these networks constitute a unique and freely accessible framework within which to generate mechanistic hypotheses around the transcriptional interface between human signaling pathways and CoV infection.

## Results

### Generation of the CoV consensomes

We first set out to generate a set of consensomes^10^ ranking human genes based on statistical measures of the frequency of their significant differential expression in response to infection by MERS, SARS1 and SARS2 CoVs. To do this we searched the Gene Expression Omnibus (GEO) and ArrayExpress databases to identify datasets involving infection of human cells by these strains. Many of these datasets emerged from a broad-scale systematic multi-omics Pacific Northwest National Library analysis of the host cellular response to infection across a broad range of pathogens^11^. Since an important question in the development of CoV therapeutics is the extent to which CoVs have common transcriptional impacts on human cell signaling that are distinct from those of other viruses, we also searched for transcriptomic datasets involving infection by human influenza A virus (IAV). From this initial collection of datasets, we next carried out a three step quality control check as previously described^10^, yielding a total of 3.3 million data points in 156 experiments from 38 independent viral infection transcriptomic datasets (figshare File F1, section 1). Using these curated datasets, we next used consensome analysis (see Methods and previous SPP publication^10^) to generate consensomes for each CoV strain. figshare File F1 contains the full human SARS1 (Section 2), SARS2 (Section 3), MERS (Section 4) and IAV (Section 5) infection transcriptomic consensomes. To assist researchers in inferring CoV infection-associated signaling networks, the consensomes are annotated using the previously described SPP convention^10^ to indicate the identity of a gene as encoding a receptor, protein ligand, enzyme, transcription factor, ion channel or co-node (figshare File F1, sections 2-5, columns A-C).

### Ranking of interferon-stimulated genes (ISGs) in the CoV consensomes

As an initial benchmark for our CoV consensome analysis, we assembled a list of 20 canonical interferon-stimulated genes (ISGs), whose role in the anti-viral response is best characterized in the context of IAV infection^12^. As shown in Figure 1, many ISGs were assigned elevated rankings across the four viral consensomes. The mean percentile of the ISGs was however appreciably higher in the IAV (98.7^th^ percentile) and SARS1 (98.5^th^ percentile; *p* = 6e-1, t-test IAV vs SARS1) consensomes than in the SARS2 (92^nd^ percentile, *p* = 5e-2, t-test IAV v SARS2) and MERS (82^nd^ percentile; *p* = 7e-5, t-test IAV v MERS) consensomes. This is consistent with previous reports of an appreciable divergence between the IAV and SARS2 transcriptional responses with respect to the interferon response^8^. Other genes with known critical roles in the response to viral infection have high rankings in the CoV consensomes, including *NCOA7*^1^ (percentiles: SARS1, 98^th^; SARS2, 97^th^; MERS, 89^th^; IAV, 99^th^), *STAT1™* (percentiles: SARS1, 99^th^; SARS2, 98^th^; MERS, 89^th^; IAV, 99^th^) and *TAP1*^15^ (percentiles: SARS1, 99^th^; SARS2, 94^th^; MERS, 83^rd^; IAV, 99^th^). In addition to the appropriate elevated rankings for these known viral response effectors, the CoV consensomes assign similarly elevated rankings to transcripts that are largely or completely uncharacterized in the context of viral infection. Examples of such genes include *PSMB9*, encoding a proteasome 20S subunit (percentiles: SARS1, 98^th^; SARS2, 97^th^; MERS, 98^th^; IAV, 98^th^); *CSRNP1*, encoding a cysteine and serine rich nuclear protein (percentiles: SARS1, 99^th^; SARS2, 94^th^; MERS, 98^th^; IAV, 94^th^); and *CCNL1*, encoding a member of the cell cycle-regulatory cyclin family (percentiles: SARS1, 99^th^; SARS2, 94^th^; MERS, 99^th^; IAV, 97^th^). Finally, a CRISPR/Cas9 study posted as a preprint while this manuscript was under review validated 27 human genes as critical modulators of the host response to SARS2 infection of human cells^16^. Corroborating our analysis, 16 of these genes have significant (*q* < 0.05) rankings in the SARS2 consensome, including *ACE2* and *DYRK1A* (both 97^th^ percentile), *CTSL* (96^th^ percentile), *KDM6A, ATRX, PIAS1* (all 94^th^ percentile), *RAD54L2* and *SMAD3* (90^th^ percentile).

**Figure 1.**
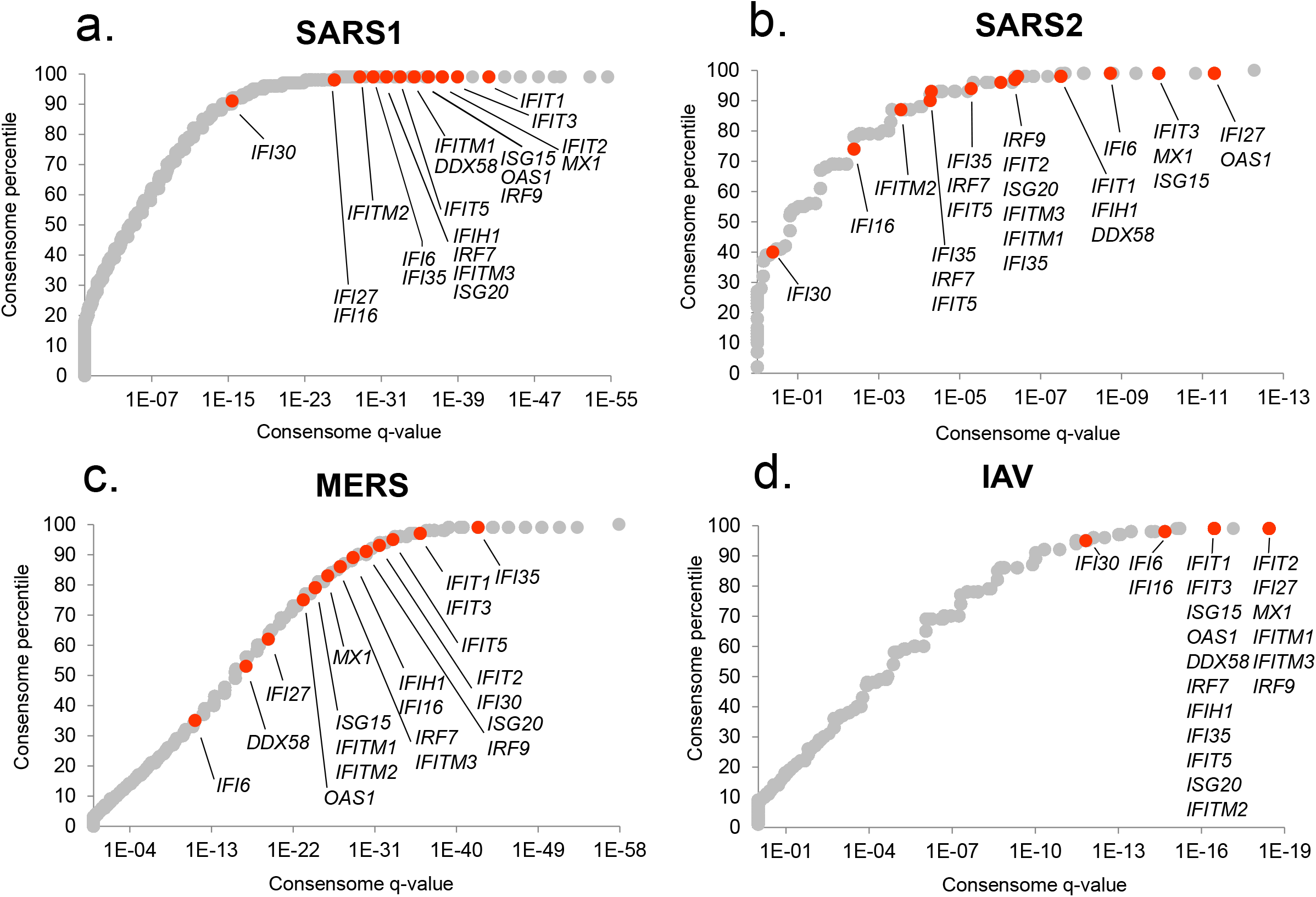
Rankings of canonical interferon-stimulated genes (ISGs) in the viral consensomes. Shown are the percentile rankings of 20 ISGS^12^ in the SARS1 **(a)**, SARS2 **(b)**, MERS **(c)** and IAV **(d)** consensomes. Note that numerous genes have identical q-value and percentile values and are therefore superimposed in the plots. Full underlying data are provided in figshare File 1. Please refer to the Methods section for a full description of the consensome algorithm.

To illuminate human signaling pathways orchestrating the transcriptional response to CoV infection, we next compared transcripts with elevated rankings in the CoV consensomes with those that have predicted high confidence regulatory relationships with cellular signaling pathway nodes. We generated four lists of genes corresponding to the MERS, SARS1, SARS2 and IAV transcriptomic consensome 95^th^ percentiles. We then retrieved genes in the 95^th^ percentiles of available SPP human transcriptomic (n = 25) consensomes and ChIP-Seq (n = 864) pathway node consensomes^10^. For convenience we will refer from hereon to genes in the 95^th^ percentile of a viral infection, node (ChIP-Seq) or node family (transcriptomic) consensome as high confidence transcriptional targets (HCTs). We then used the R GeneOverlap package^17^ to compute the extent and significance of intersections between CoV HCTs and those of the pathway nodes or node families. We interpreted the extent and significance of intersections between HCTs for CoVs and pathway node or node families as evidence for a biological relationship between loss or gain of function of that node (or node family) and the transcriptional response to infection by a specific virus.

Results of viral infection and signaling node HCT intersection analyses are shown in Figure 2 (based on receptor and enzyme family transcriptomic consensomes), Figures 3 and 4 (based on ChIP-Seq consensomes for transcription factors and enzymes, respectively) and figshare File F2 (based on ChIP-Seq consensomes for selected co-nodes). figshare File F1, sections 6 (node family transcriptomic HCT intersection analysis) and 7 (node ChIP-Seq HCT intersection analysis) contain the full underlying numerical data. We surveyed *q* < 0.05 HCT intersections to identify (i) canonical inflammatory signaling pathway nodes with characterized roles in the response to CoV infection, thereby validating the consensome approach in this context; and (ii) evidence for nodes whose role in the transcriptional biology of CoV infection is previously uncharacterized, but consistent with their roles in the response to other viral infections. In the following sections all *q*-values refer to those obtained using the GeneOverlap analysis package in R^17^.

**Figure 2.**
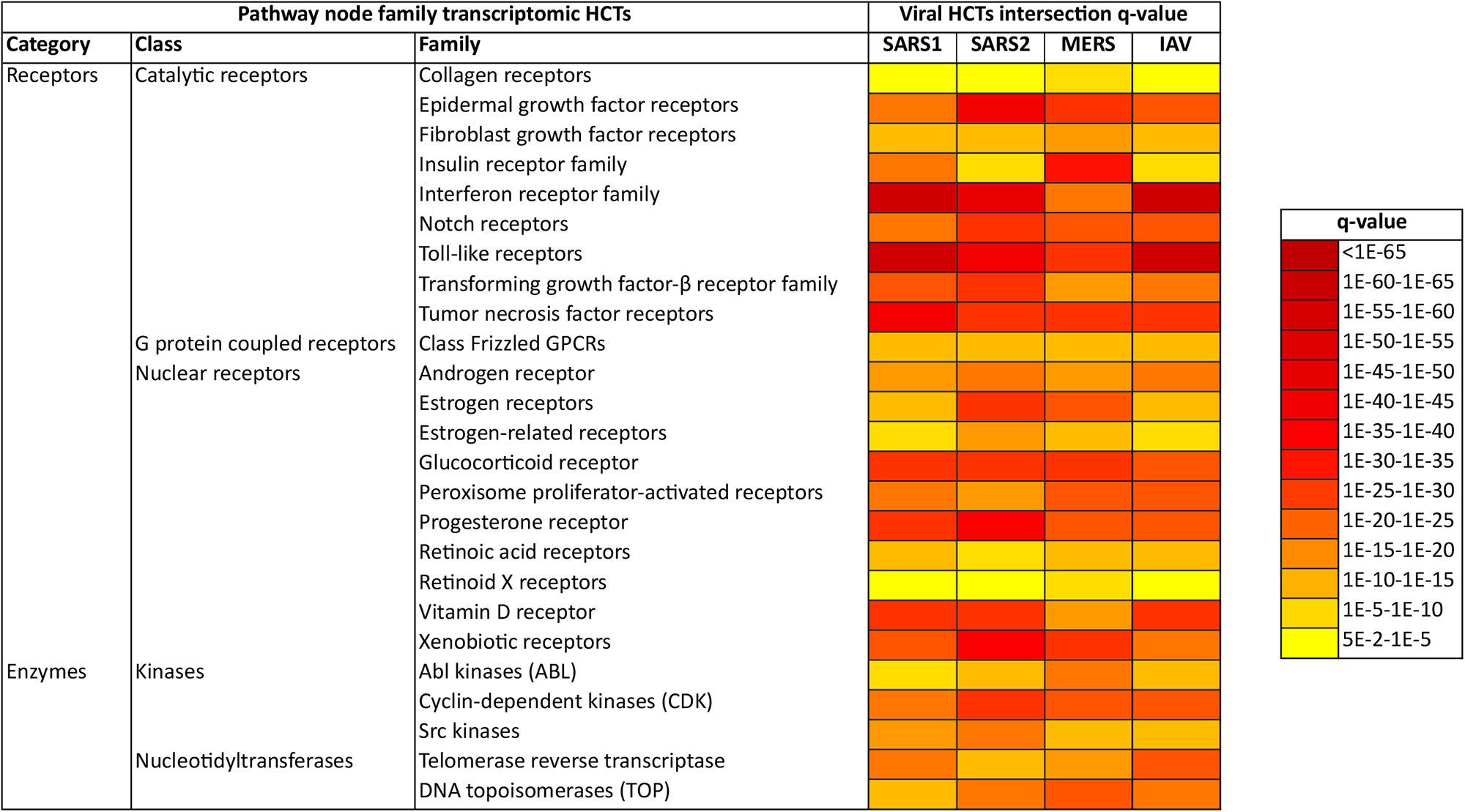
High confidence transcriptional target (HCT) intersection analysis of viral infection and human receptors or signaling enzymes based on transcriptomic consensomes. Full numerical data are provided in figshare File F1, section 6. Due to space constraints some class and family names may differ slightly from those in the SPP knowledgebase. All q-values refer to those obtained using the GeneOverlap analysis package in R^17^.

**Figure 3.**
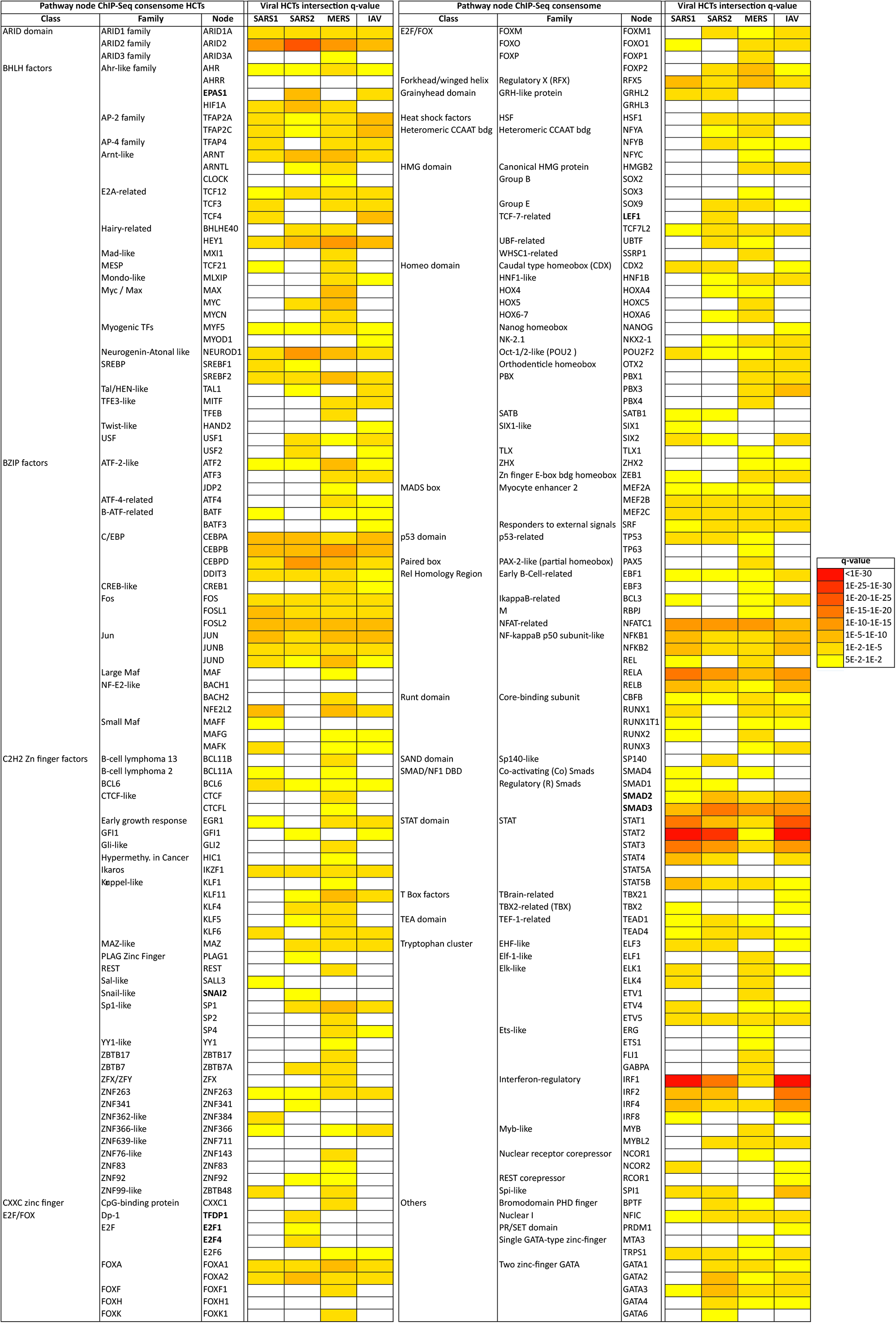
High confidence transcriptional target (HCT) intersection analysis of viral infection and human transcription factors based on ChIP-Seq consensomes. White cells represent *q* > 5e-2 intersections. Full numerical data are provided in figshare File F1, section 7. Due to space constraints some class and family names may differ slightly from those in the SPP knowledgebase. All q-values refer to those obtained using the GeneOverlap analysis package in R^17^.

**Figure 4.**
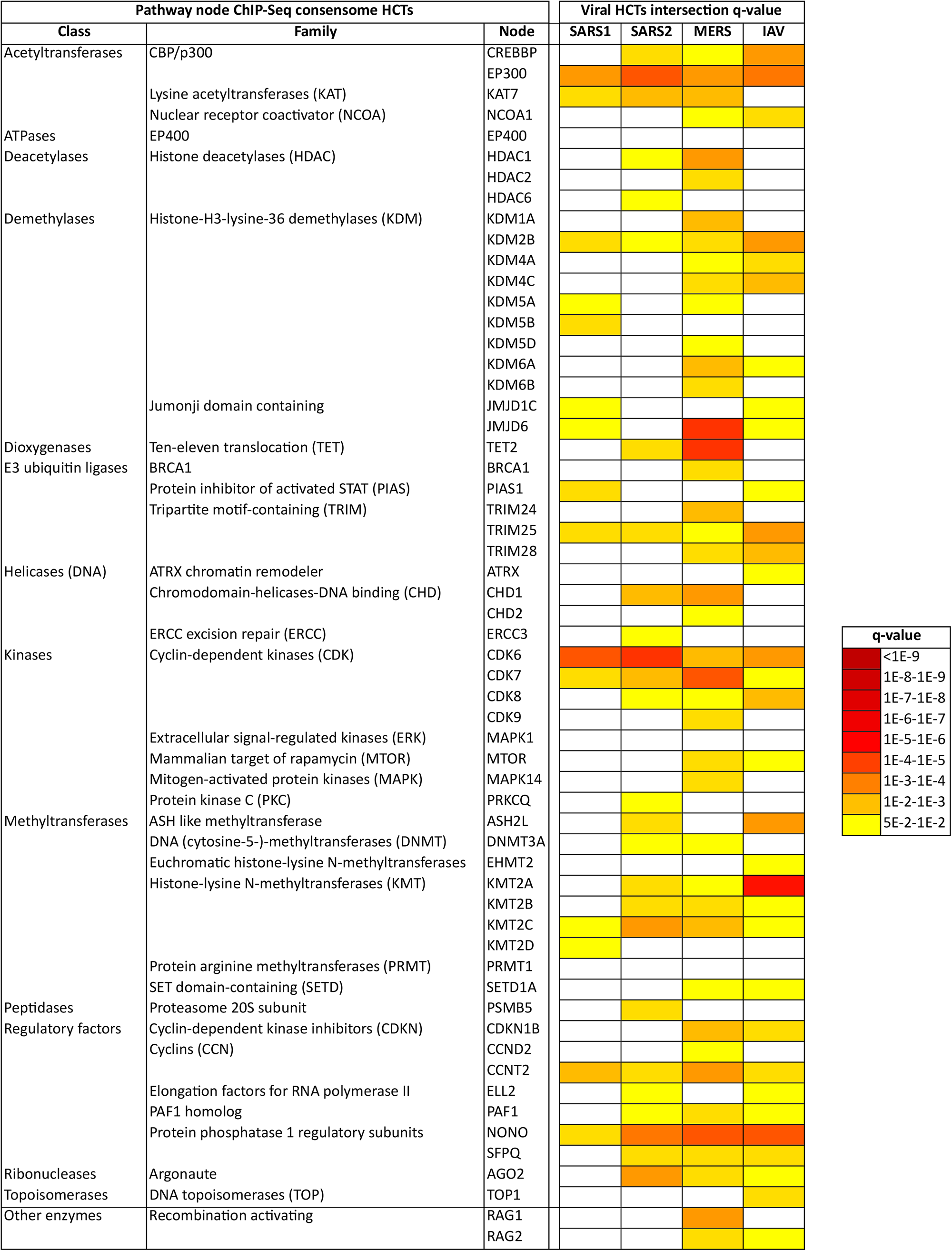
High confidence transcriptional target (HCT) intersection analysis of viral infection and human signaling enzymes based on ChIP-Seq consensomes. White cells represent non-significant (*q* > 5e-2) intersections. Full numerical data are provided in figshare File F1, section 7. Due to space constraints some class and family names may differ slightly from those in the SPP knowledgebase. All q-values refer to those obtained using the GeneOverlap analysis package in R^17^.

#### Receptors

Reflecting their well-documented roles in the response to CoV infection^18–21^, we observed appreciable significant intersections between CoV HCTs and those of the toll-like (TLRs; q-values: SARS1, 3e-85; SARS2, 5e-49; MERS, 2e-33), interferon (IFNR; q-values: SARS1, 1e-109; SARS2, 6e-53; MERS, 1e-24) and tumor necrosis factor (TNFR; q-values: SARS1, 1e-48; SARS2, 1e-35; MERS, 5e-32) receptor families (Fig. 2). HCT intersections between CoV infection and receptor systems with previously uncharacterized connections to CoV infection, including epidermal growth factor receptors (EGFR; q-values: SARS1, 4e-21; SARS2, 3e-48; MERS, 1e-35), and Notch receptor signaling (q-values: SARS1, 6e-24; SARS2, 2e-33; MERS, 2e-29; Fig. 2), are consistent with their known role in the context of other viral infections^22–26^. The Notch receptor HCT intersection points to a possible mechanistic basis for the potential of Notch pathway modulation in the treatment of SARS2^27^. The strong HCT intersection between CoV infection and xenobiotic receptors (q-values: SARS1, 1e-30; SARS2, 1e-44; MERS, 5e-32; Fig. 2) reflects work describing a role for pregnane X receptor in innate immunity^28^ and points to a potential role for members of this family in the response to CoV infection. In addition, the robust intersection between HCTs for SARS2 infection and vitamin D receptor (*q* = 2e-35) is interesting in light of epidemiological studies suggesting a link between risk of SARS2 infection and vitamin D deficiency^29,30^. Consistent with a robust signature for the glucocorticoid receptor across all CoVs (GR; q-values: SARS1, 3e-35; SARS2, 1e-35; MERS, 7e-32), while this paper was under review, studies were published showing the GR agonist dexamethasone was a successful therapeutic for SARS2 infection^31^. Finally, and also while this paper was under review, in vitro analyses confirmed our predictions of the modulation by SARS2 infection of ErbB/EGFR^20,32^ and TGFBR^16,32^ signaling systems (Fig. 2).

#### Transcription factors

Not unexpectedly – and speaking again to validation of the consensomes - the strongest and most significant CoV HCT intersections were observed for HCTs for known transcription factor mediators of the transcriptional response to CoV infection, including members of the NFκB (q-value ranges: SARS1, 1e-7-1e-9; SARS2, 9e-3-2e-3; MERS, 1e-3-1e-4)^33–35^, IRF (q-value ranges: SARS1, 2e-2-1e-31; SARS2, 2e-4-1e-17; MERS, 9e-4-7e-5)^36^ and STAT (q-value ranges: SARS1, 1e-7-1e-55; SARS2, 2e-3-3e-29; MERS, 5e-2-3e-5)^37–39^ transcription factor families (Fig. 3). Consistent with the similarity between SARS1 and IAV consensomes with respect to elevated rankings of ISGs (Fig. 2a & d), the IRF1 HCT intersection was strongest with the SARS1 (*q* = 2e-34) and IAV (*q* = 3e-49) HCTs. Corroborating our finding of a strong intersection between STAT2 and SARS2 infection HCTs (*q =* 3e-29), a study that appeared while this manuscript was under review showed that STAT2 plays a prominent role in the response to SARS2 infection of Syrian hamsters^40^. HCT intersections for nodes originally characterized as having a general role in RNA Pol II transcription, including TBP (q-values: SARS1, 2e-10; SARS2, 6e-23; MERS, 3e-16), GTF2B/TFIIB (q-values: SARS1, 7e-10; SARS2, 3e-23; MERS, 9e-14) and GTF2F1 (q-values: SARS1, 2e-4; SARS2, 2e-13; MERS, 5e-5) were strong across all CoVs, and particularly noteworthy in the case of SARS2. In the case of GTF2B, these data are consistent with previous evidence identifying it as a specific target for orthomyxovirus^41^, and the herpes simplex^42^ and hepatitis B^43^ viruses. Moreover, a proteomic analysis that appeared in BioRXiv while this paper was under review identified a high confidence interaction between GTF2F2 and the SARS2 NSP9 replicase^32^.

In general, intersections between viral infection and ChIP-Seq enrichments for transcription factors and other nodes were more specific for individual CoV infection HCTs (compare Fig. 2 with Figs. 3 & 4 and figshare File F1, sections 6 and 7). This is likely due to the fact that ChIP-Seq consensomes are based on direct promoter binding by a specific node antigen, whereas transcriptomic consensomes encompass both direct and indirect targets of specific receptor and enzyme node families.

#### Enzymes

Compared to the roles of receptors and transcription factors in the response to viral infection, the roles of signaling enzymes are less well illuminated – indeed, in the context of CoV infection, they are entirely unstudied. Through their regulation of cell cycle transitions, cyclin-dependent kinases (CDKs) play important roles in the orchestration of DNA replication and cell division, processes that are critical in the viral life cycle. CDK6, which has been suggested to be a critical G1 phase kinase^44,45^, has been shown to be targeted by a number of viral infections, including Kaposi’s sarcoma-associated herpesvirus^46^ and HIV-1^47^. Consistent with this common role across distinct viral infections, we observed robust intersection between the CDK family HCTs (q-values: SARS1, 8e-23; SARS2, 2e-31; MERS, 1e-30; Fig. 2) and the CDK6 HCTs (q-values: SARS1, 1e-7; SARS2, 8e-8; MERS, 3e-4; Fig. 4) and those of all viral HCTs. As with the TLRs, IFNRs and TNFRs, which are known to signal through CDK6^48–50^, intersection with the CDK6 HCTs was particularly strong in the case of the SARS2 HCTs (Fig. 4). Again, the subsequent proteomic analysis we alluded to earlier^32^ independently corroborated our prediction of a role for CDK6 in the response to SARS2 infection.

CCNT2 is another member of the cyclin family that, along with CDK9, is a component of the viral-targeted p-TEFB complex^51^. Reflecting a potential general role in viral infection, appreciable intersections were observed between the CCNT2 HCTs and all viral HCTs (q-values: SARS1,4e-4; SARS2, 6e-3; MERS, 7e-5; Fig. 4). Finally in the context of enzymes, the DNA topoisomerases have been shown to be required for efficient replication of simian virus 40^52^ and Ebola^53^ viruses. The prominent intersections between DNA topoisomerase-dependent HCTs and the CoV HCTs (q-values: SARS1, 3e-15; SARS2, 6e-21; MERS, 1e-26; Fig. 4) suggest that it may play a similar role in facilitating the replication of these CoVs.

### Hypothesis generation use cases

We next wished to show how the CoV consensomes and HCT intersection networks, supported by existing canonical literature knowledge, enable the user to generate novel hypotheses around the transcriptional interface between CoV infection and human cellular signaling pathways. Given the current interest in SARS2, we have focused our use cases on that virus. In addition to these use cases, figshare File F2 contains a number of additional use cases omitted from the main text due to space constraints. Unless otherwise stated, all *q*-values below were obtained using the GeneOverlap analysis package in R^17^. We stress that all use cases represent preliminary *in silico* evidence only, and require rigorous pressure-testing at the bench for full validation.

### Hypothesis generation use case 1: transcriptional regulation of the SARS2 receptor gene, *ACE2*

*ACE2*, encoding membrane-bound angiotensin converting enzyme 2, has gained prominence as the target for cellular entry by SARS1^54^ and SARS2^55^. An important component in the development of ACE2-centric therapeutic responses is an understanding of its transcriptional responsiveness to CoV infection. Interestingly, based on our CoV consensome analysis, *ACE2* is more consistently transcriptionally responsive to infection by SARS CoVs (SARS1: 98^th^ percentile, consensome *q* value (CQV)^10^ *=* 1e-25; SARS2: 97^th^ percentile, CQV *=* 4e-7) than by IAV (78^th^ percentile, CQV *=* 3e-8) or MERS (49^th^ percentile, CQV *=* 2e-16; figshare File F1, sections 2-5). The data points underlying the CoV consensomes indicate evidence for tissue-specific differences in the nature of the regulatory relationship between *ACE2* and viral infection. In response to SARS1 infection, for example, *ACE2* is induced in pulmonary cells but repressed in kidney cells (Fig. 5). On the other hand, in response to SARS2 infection, *ACE2* is repressed in pulmonary cells - a finding corroborated by other studies ^56,57^ - but inducible in gastrointestinal cells (Fig. 5). These data may relate to the selective transcriptional response of *ACE2* to signaling by IFNRs (92^nd^ percentile; figshare File F1, section 8) rather than TLRs (48^th^ percentile; figshare File F1, section 9) or TNFRs (13^th^ percentile, figshare File F1, section 10). While this manuscript was under review, another study appeared confirming repression of induction of *ACE2* by interferon stimulation and by IAV infection^58^. Our data reflect a complex transcriptional relationship between *ACE2* and viral infection that may be illuminated in part by future single cell RNA-Seq analysis in the context of clinical or animal models of SARS2 infection.

**Figure 5.**
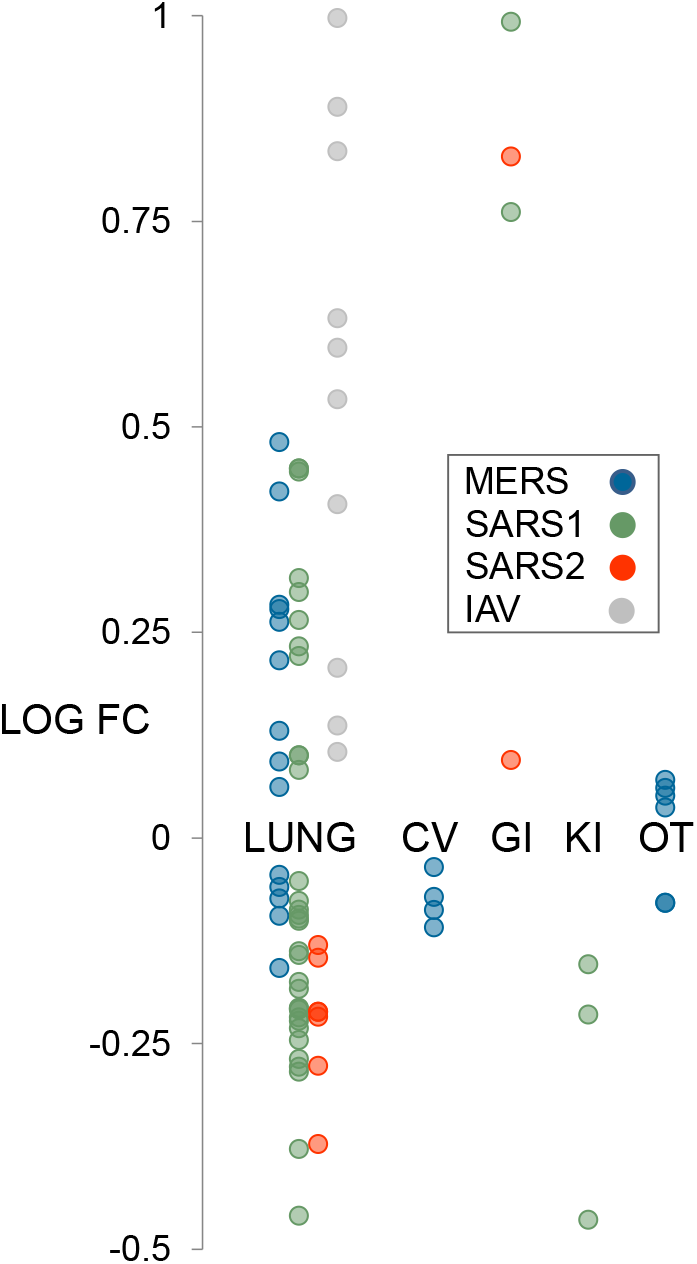
Hypothesis generation use case 1: strain- and tissue-specific regulation of *ACE2* in response to CoV infection of human cells. All data points are p < 0.05. Refer to figshare File F1, section 1 for full details on the underlying datasets.

### Hypothesis generation use case 2: evidence for antagonistic cross-talk between progesterone receptor and interferon receptor signaling in the airway epithelium

A lack of clinical data has so far prevented a definitive evaluation of the connection between pregnancy and susceptibility to SARS2 infection in CoVID-19. That said, SARS2 infection is associated with an increased incidence of pre-term deliveries^59^, and pregnancy has been previously associated with the incidence of viral infectious diseases, particularly respiratory infections^60,61^. We were therefore interested to observe consistent intersections between the progesterone receptor (PGR) HCTs and CoV infection HCTs (q-values: SARS1, 3e-35; SARS2, 5e-41; MERS 5e-28), with the intersection being particularly evident in the case of the SARS2 HCTs (Fig. 2; figshare File F1, section 6). To investigate the specific nature of the crosstalk implied by this transcriptional intersection in the context of the airway epithelium, we first identified a set of 12 genes that were HCTs for both SARS2 infection and PGR. Interestingly, many of these genes encode members of the classic interferon-stimulated gene (ISG) response pathway^12^. We then retrieved two SPP experiments involving treatment of A549 airway epithelial cells with the PGR full antagonist RU486 (RU), alone or in combination with the GR agonist dexamethasone (DEX). As shown in Figure 6, there was unanimous correlation in the direction of regulation of all 12 genes in response to CoV infection and PGR loss of function. These data are consistent with the reported pro-inflammatory effects of RU486 in a mouse model of allergic pulmonary inflammation^62^. Interestingly, SARS2-infected pregnant women are often asymptomatic^63,64^. Based on our data, it can be reasonably hypothesized that suppression of the interferon response to SARS2 infection by elevated circulating progesterone during pregnancy may contribute to the asymptomatic clinical course. Indeed, crosstalk between progesterone and inflammatory signaling is well characterized in the reproductive system, most notably in the establishment of uterine receptivity^65^ as well as in ovulation^66^. Consistent with our hypothesis, while this paper was under review, a clinical trial was launched to evaluate the potential of progesterone for treatment of COVID-19 in hospitalized men^67^. Interestingly, and also while this paper was under review, a paper appeared showing that progesterone inhibited SARS2 replication in African green monkey kidney Vero 6 cells^68^. These results indicate an additional mechanism, distinct from its potential crosstalk with the interferon response, by which progesterone signaling may impact SARS2 infection.

**Figure 6.**
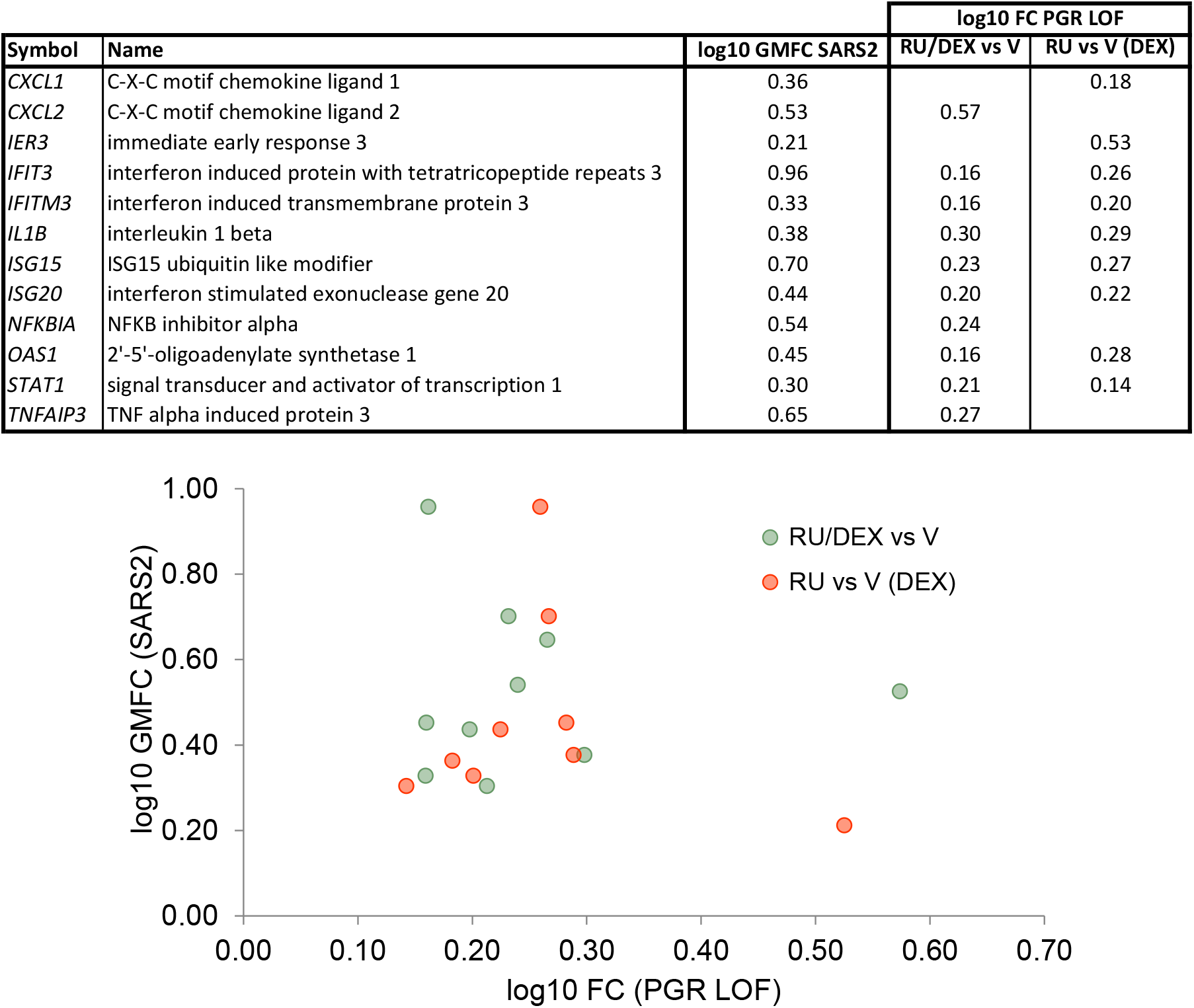
Hypothesis generation use case 2: antagonism between PGR and SARS2 inflammatory signaling in the regulation of viral response genes in the airway epithelium. GMFC: geometric mean fold change. PGR loss of function experiments were retrieved from the SPP knowledgebase^128^.

### Hypothesis generation use case 3: association of an epithelial to mesenchymal transition transcriptional signature with SARS2 infection

Epithelial to mesenchymal transition (EMT) is the process by which epithelial cells lose their polarity and adhesive properties and acquire the migratory and invasive characteristics of mesenchymal stem cells^69^. EMT is known to contribute to pulmonary fibrosis^70^, acute interstitial pneumonia^71^ and acute respiratory distress syndrome (ARDs)^72^, all of which have been reported in connection with SARS2 infection in COVID-19. We were interested to note therefore that significant HCT intersections for three well characterized EMT-promoting transcription factors were specific to SARS2 infection (q-values: SNAI2/Slug^76^, 2e-2; EPASI/HIF2α^77^, 9e-9; LEF1^78^, 1e-3; Fig. 3, bold symbols; figshare File F1, section 7). Consistent with this, intersections between HCTs for TGFBRs, SMAD2 and SMAD3, known regulators of EMT transcriptional programs^79^ – were stronger with HCTs for SARS2 (q-values: TGFBRs, 2e-31; SMAD2, 2e-7; SMAD3, 5e-17) than with those of SARS1 (q-values: TGFBRs, 6e-29; SMAD2, 2e-2; SMAD3, 3e-9) and MERS (q-values: TGFBRs, 1e-16; SMAD2, 3e-3; SMAD3, 2e-12) – see also Figs. 2 and 3 and figshare File F1, sections 6 and 7). Moreover, a recent CRISPR/Cas9 screen identified a requirement for both TGFBR signaling and *SMAD3* in mediating SARS2 infection^16^.

To investigate the connection between SARS2 infection and EMT implied by these HCT intersections, we then computed intersections between the individual viral HCTs and a list of 335 genes manually curated from the research literature as EMT markers (figshare File F1, section 11). In agreement with the HCT intersection analysis, we observed significant enrichment of members of this gene set within the SARS2 HCTs (*q =* 4e-14), but not the SARS1 or MERS (both *q =* 2e-1) HCTs (Fig. 7a). Consistent with previous reports of a potential link between EMT and IAV infection^81^, we observed significant intersection between the EMT signature and the IAV HCTs (*q* = 1e-04).

**Figure 7.**
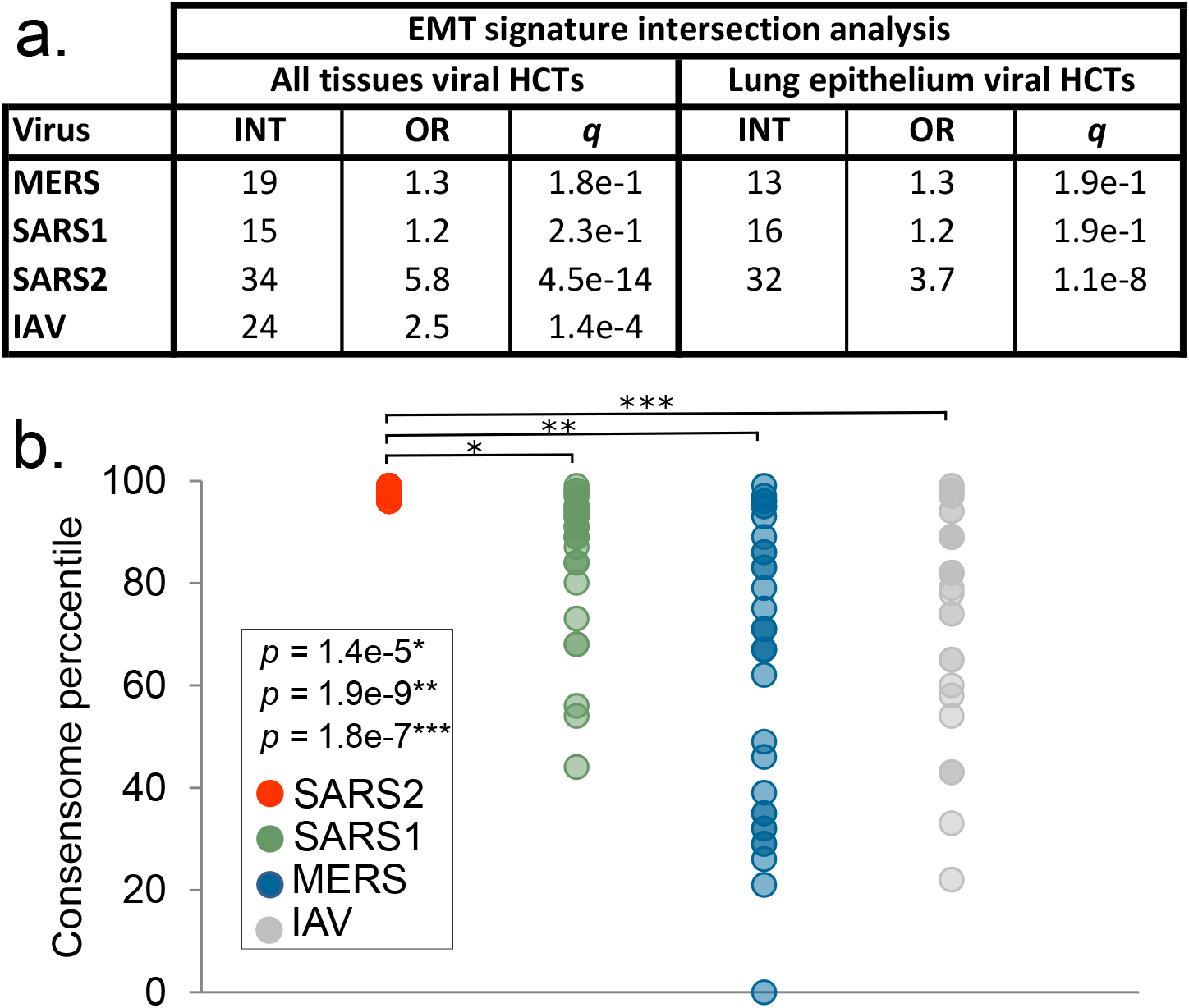
Hypothesis generation use case 3: evidence for a SARS2 infection-associated EMT transcriptional signature. **a.** CoV HCT intersection with the literature-curated EMT signature for all-biosample and lung epithelium-specific consensomes. The IAV consensome is comprised of lung epithelial cell lines and was therefore omitted from the lung epithelium-only consensome analysis. Refer to the column “EMT” in figshare File F1, section 3 for the list of EMT SARS2 HCTs. q-values refer to those obtained using the GeneOverlap analysis package in R^17^. **b.** Comparison of mean percentile ranking of the EMT-associated SARS2 HCTs across viral consensomes. Note that SARS2 HCTs are all in the 97-99^th^ percentile and are therefore superimposed in the scatterplot. Indicated are the results of the two-tailed two sample t-test assuming equal variance comparing the percentile rankings of the SARS2 EMT HCTs across the four viral consensomes.

One possible explanation for the selective intersection between the literature EMT signature and the SARS2 HCTs relative to SARS1 and MERS was the fact that the SARS2 consensome was exclusively comprised of epithelial cell lines, whereas the SARS1 and MERS consensomes included non-epithelial cell biosamples (figshare File F1, section 1). To exclude this possibility therefore, we next calculated airway epithelial cell-specific consensomes for SARS1, SARS2 and MERS and computed intersections between their HCTs and the EMT signature. We found that significant intersection of the EMT signature with the CoV HCTs remained specific to SARS2 (q-values: SARS1, 2e-1; SARS2, 1e-8; MERS, 2e-1) in the lung epithelium-specific CoV consensomes.

We next retrieved the canonical EMT genes in the SARS2 HCTs and compared their percentile rankings with the other CoV consensomes. Although some EMT genes, such as *CXCL2* and *IRF9*, had elevated rankings across all four viral consensomes, the collective EMT gene signature had a significantly higher mean percentile value in the SARS2 consensome than in each of the other viral consensomes (Fig. 7b; SARS2 mean percentile = 97.5; SARS1 mean percentile = 86, *p* = 1e-5, t-test; MERS mean percentile = 63, *p* = 1e-9, t-test; IAV mean percentile = 76, p = 2e-7, t-test). A column named “EMT” in figshare File F1, sections 2 (SARS1), 3 (SARS2), 4 (MERS) and 5 (IAV) identifies the ranking of the EMT genes in each of the viral consensomes.

Given that EMT has been linked to ARDs^72^, we speculated that the evidence connecting EMT and SARS2 acquired through our analysis might be reflected in the relatively strong intersection between ARDs markers in SARS2 HCTs compared to other viral HCTs. To test this hypothesis we carried out a PubMed search to identify a set of 88 expression biomarkers of ARDs or its associated pathology, acute lung injury (ALI). A column named “ALI/ARDs” in figshare File F1, sections 2 (SARS1), 3 (SARS2) 4 (MERS) and 5 (IAV) identifies the expression biomarker genes using the PubMed identifiers for the original studies in which they were identified. Consistent with our hypothesis, we observed appreciable intersections between this gene set and the HCTs of all four viruses (SARS1 odds ratio (OR) = 7, *q* = 5e-9; SARS2 OR = 10.4, *q* = 1e-9; MERS, OR = 4.2, *q* = 2e-5; IAV OR = 6.8; *q* = 9e-8) with a particularly strong intersection evident in the SARS2 HCTs.

Although EMT has been associated with infection by transmissible gastroenteritis virus^82^ and IAV^81^, this is to our knowledge the first evidence connecting CoV infection, and specifically SARS2 infection, to an EMT signature. Interestingly, lipotoxin A4 has been shown to attenuate lipopolysaccharide-induced lung injury by reducing EMT^83^. Moreover, several members of the group of SARS2-induced EMT genes have been associated with signature pulmonary comorbidities of CoV infection, including *ADAR*^84^, *CLDN1^85^* and *SOD2*^86^. Of note in the context of these data is the fact that signaling through two SARS2 cellular receptors, ACE2/AT2 and CD147/basigin, has been linked to EMT in the context of organ fibrosis Finally, while this manuscript was under review, a preprint was posted that described EMT-like transcriptional and metabolic changes in response to SARS2 infection^90^. Collectively, our data indicate that EMT warrants further investigation as a SARS2-specific pathological mechanism.

### Hypothesis generation use case 4: SARS2 repression of E2F family HCTs encoding cell cycle regulators

Aside from EPAS1 and SNAI2, the only other transcription factors with significant HCT intersections that were specific to the SARS2 HCTs were the E2F/FOX class members E2F1 (q-values: SARS1, 5e-1; SARS2, 1e-2; MERS, 4e-1), E2F3 (q-values: SARS1, 6e-1; SARS2, 5e-2; MERS, 7e-1), E2F4 (q-values: SARS1, 1; SARS2, 9e-3; MERS, 1) and TFDP1/Dp-1 (q-values: SARS1, 1; SARS2, 3e-4; MERS, 1; Fig. 3, bold symbols; figshare File F1, section 7). These factors play well-documented interdependent roles in the promotion (E2F1, E2F3, TFDP1) and repression (E2F4) of cell cycle genes^91,92^. Moreover, E2F family members are targets of signaling through EGFRs^93^ and CDK6^94^, both of whose HCTs had SARS2 HCT intersections that were stronger those of the other CoVs (EGFRs: q-values: SARS1, 4e-21; SARS2, 3e-48; MERS, 1e-35; CDK6: q-values: SARS1, 1e-7; SARS2, 8e-8; MERS, 2e-4); Figs. 2 & 4). Based on these data, we speculated that SARS2 infection might impact the expression of E2F-regulated cell cycle genes more efficiently than other CoVs. To investigate this we retrieved a set of SARS2 HCTs that were also HCTs for at least three of E2F1, E2F3, E2F4 and TFDP1 (figshare File F1, section 3, columns P-T). Consistent with the role of E2F/Dp-1 nodes in the regulation of the cell cycle, many of these genes – notably *CDK1, PCNA, CDC6, CENPF* and *NUSAP1* – are critical positive regulators of DNA replication and cell cycle progression^95–99^ and are known to be transcriptionally induced by E2Fs^100–103^. Strikingly, with the exception of *E2F3*, all were consistently repressed in response to SARS2 infection (Fig. 8a). To gain insight into the relative efficiency with which the four viruses impacted expression of the E2F/Dp-1 HCT signature, we compared their mean percentile values across the viral consensomes. Consistent with efficient repression of the E2F/Dp-1 HCTs by SARS2 infection relative to other viruses, their mean percentile ranking was appreciably higher in the SARS2 consensome (97^th^ percentile) than in the SARS1 (76^th^ percentile; *p* = 6e-12, t-test), MERS (71.2 percentile; p = 9e-6, t-test) and IAV (71.2 percentile; p = 2e-5, t-test) consensomes (Fig. 8b). Although manipulation of the host cell cycle and evasion of detection through deregulation of cell cycle checkpoints has been described for other viruses^104–106^, this represents the first evidence for the profound impact of SARS2 infection on host cell cycle regulatory genes, potentially through disruption of E2F mediated signaling pathways. The SARS2 infection-mediated induction of *E2F3* (Fig. 8a) may represent a compensatory response to transcriptional repression of other E2F family members, as has been previously observed for this family in other contexts^107,108^. Consistent with our prediction in this use case, while this paper was in revision, a study appeared showing that infection by SARS2 results in cell cycle arrest^109^. Our results represent evidence that efficient modulation by SARS2 of E2F signaling, resulting in repression of cell cycle regulatory genes, may contribute to its unique pathological impact.

**Figure 8.**
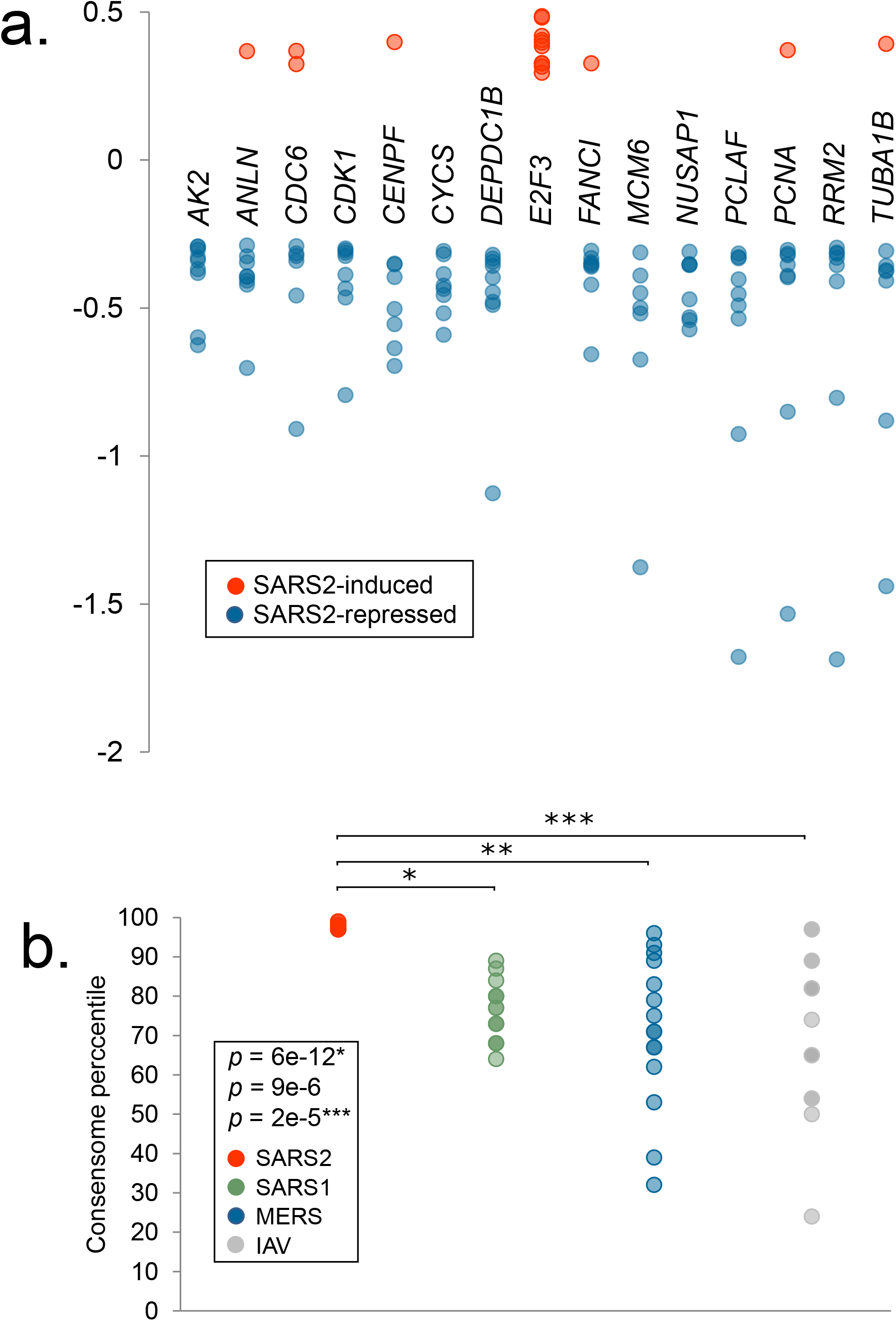
Hypothesis generation use case 4: efficient SARS2 repression of E2F family HCTs encoding key cell cycle regulators. **a.** Relative abundance of E2F HCT cell cycle regulators in response to SARS2 infection. **b.** Comparison of SARS2, SARS1, MERS and IAV consensome percentiles of the E2F HCT cell cycle regulators. Indicated are the results of the two-tailed two sample t-test assuming equal variance comparing the percentile rankings of the SARS2 EMT HCTs across the four viral consensomes.

### Visualization of the CoV transcriptional regulatory networks in the Signaling Pathways Project knowledgebase and Network Data Exchange repository

To enable researchers to routinely generate mechanistic hypotheses around the interface between CoV infection human cell signaling, we next made the consensomes and accompanying HCT intersection analyses freely available to the research community in the SPP knowledgebase and the Network Data Exchange (NDEx) repository. Table 1 contains digital object identifier (DOI)-driven links to the consensome networks in SPP and NDEx, and to the HCT intersection networks in NDEx.

**Table 1.**
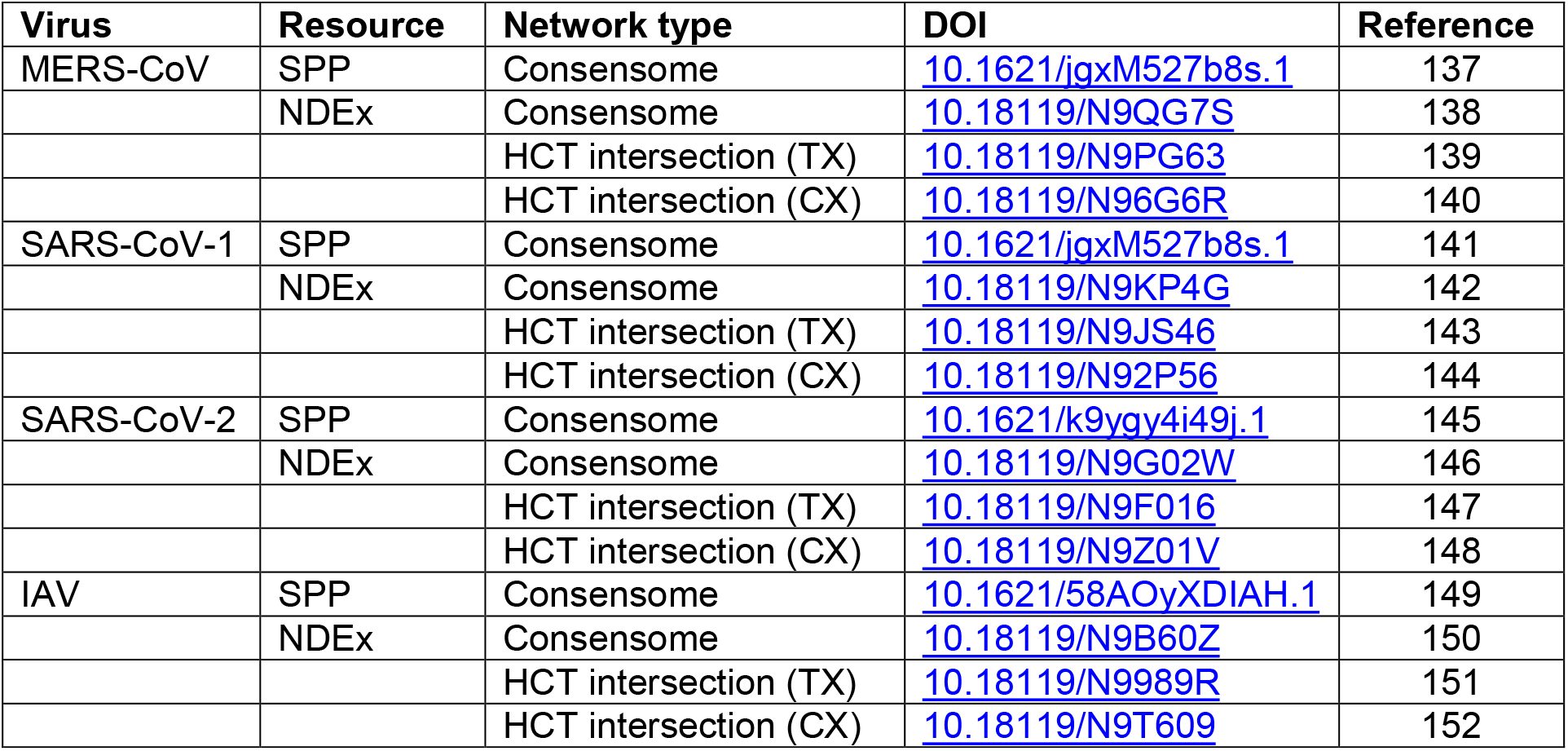
DOI-driven links to consensomes and HCT intersection networks. SPP DOIs point to the web browser version of the consensome, which contains a downloadable version of the full consensome. For clarity of visualization, NDEx consensome DOIs point to networks containing transcripts in the top 5% of each consensome (i.e. HCTs for each viral infection); the full consensome network can be reached from this page. Similarly, NDEx HCT intersection DOIs point to networks containing nodes in the top 5% of each HCT intersection network; the full HCT intersection network can be reached from this page. TX, transcriptomic node family intersection; CX, ChIP-Seq node intersection.

We have previously described the SPP biocuration pipeline, database and web application interface^10^. Figure 9 shows the strategy for consensome data mining on the SPP website. The individual CoV consensomes can be accessed by configuring the SPP Ominer query form as shown, in this example for the SARS2 consensome (Fig. 9a). Figure 9b shows the layout of the consensomes, showing gene symbol, name, percentile ranking and other essential information. Genes in the 90^th^ percentile of each consensome are accessible via the user interface, with the full consensomes available for download in a tab delimited text file. Target gene symbols in the consensome link to the SPP Regulation Report, filtered to show only experimental data points that contributed to that specific consensome (Fig. 9c). This view gives insights into the influence of tissue and cell type context on the regulatory relationship. These filtered reports can be readily converted to default Reports that show evidence for regulation of a specific gene by other signaling pathway nodes. As previously described, pop-up windows in the Report provide experimental details, in addition to links to the parent dataset (Fig. 9d), curated accordingly to our previously described protocol^10^. Per FAIR data best practice, CoV infection datasets – like all SPP datasets – are associated with detailed descriptions, assigned a DOI, and linked to the associated article to place the dataset in its original experimental context (Fig. 9d). The full list of datasets is available for browsing in the SPP Dataset listing (https://www.signalingpathways.org/index.jsf).

**Figure 9.**
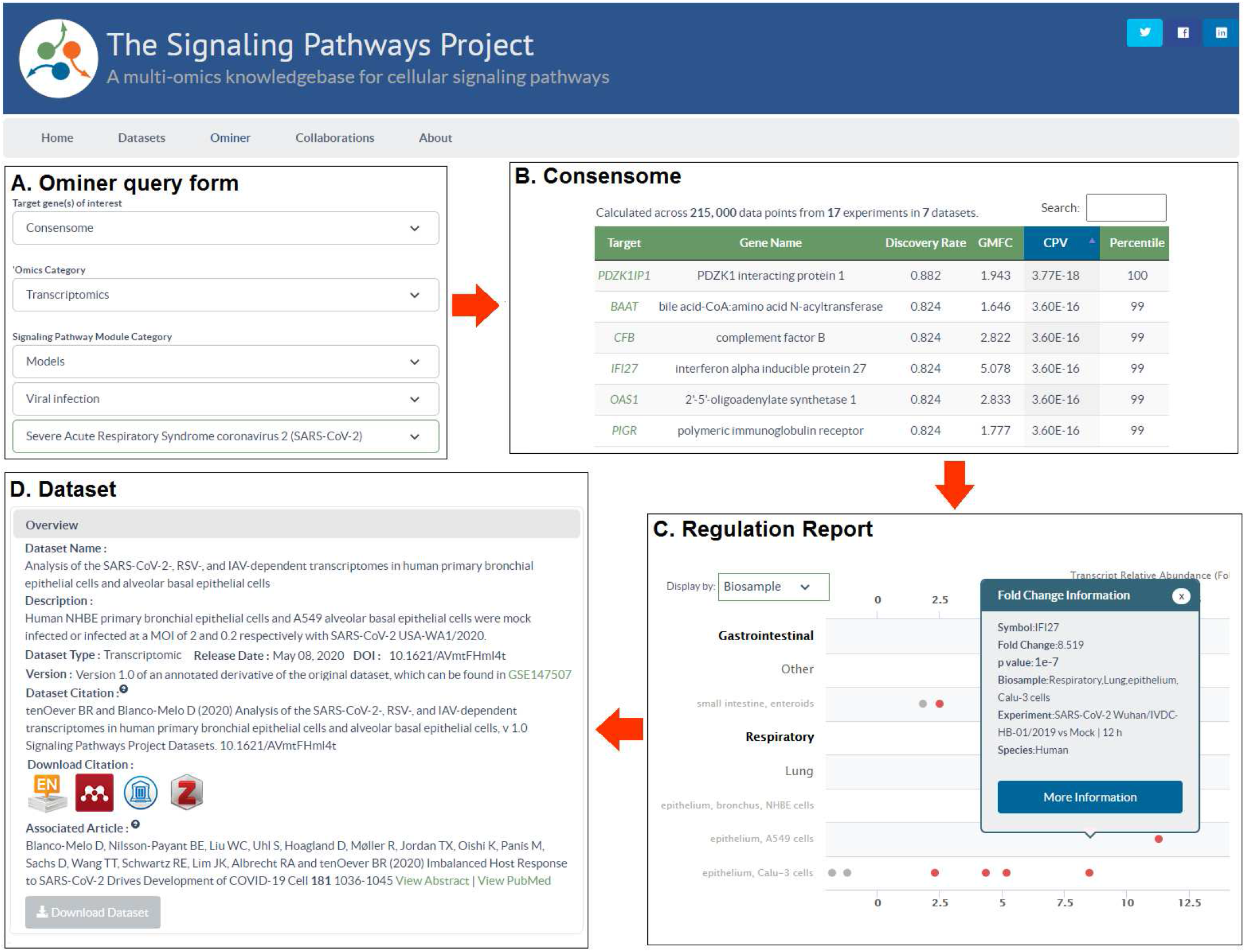
Mining of CoV consensomes and underlying data points in the SPP knowledgebase. **a.** The Ominer query form can be configured as shown to access the CoV infection consensomes. In the example shown, the user wishes to view the SARS2 consensome. **b**. Consensomes are displayed in a tabular format. Target transcript symbols in the consensomes link to SPP transcriptomic Regulation Reports (**c**) **c.** Regulation Reports for consensome transcripts are filtered to show only data points that contributed to their consensome ranking. Clicking on a data point opens a Fold Change Information window that links to the SPP curated version of the original archived dataset (d). **d.** Like all SPP datasets, CoV infection datasets are comprehensively aligned with FAIR data best practice and feature human-readable names and descriptions, a DOI, one-click addition to citation managers, and machine-readable downloadable data files. For a walk-through of CoV consensome data mining options in SPP, please refer to the accompanying YouTube video (http://tiny.cc/2i56rz).

The NDEx repository facilitates collaborative publication of biological networks, as well as visualization of these networks in web or desktop versions of the popular and intuitive Cytoscape platform^110–112^. Figure 10 shows examples of consensome and HCT intersection network visualizations within the NDEx user interface. For ease of viewing, the initial rendering of the full SARS2 (Fig. 10a) and other consensome networks shows a sample (Fig. 10a, red arrow 1) containing only the top 5% of regulated transcripts; the full data can be explored using the “Neighborhood Query” feature available at the bottom of the page (red arrow 2). The integration in NDEx of the popular Cytoscape desktop application enables any network to be seamlessly be imported in Cytoscape for additional analysis (red arrow 3). Zooming in on a subset of the SARS2 consensome (orange box) affords an appreciation of the diversity of molecular classes that are transcriptionally regulated in response to SARS2 infection (Fig. 10b). Transcript size is proportional to rank percentile, and edge weight is proportional to the transcript geometric mean fold change (GMFC) value. Selecting a transcript allows the associated consensome data, such as rank, GMFC and family, to be examined in detail using the information panel (Fig. 10b, right panel). Highlighted to exemplify this feature is IL6, an inflammatory ligand that has been previously linked to SARS2 pathology^8,113^.

**Figure 10.**
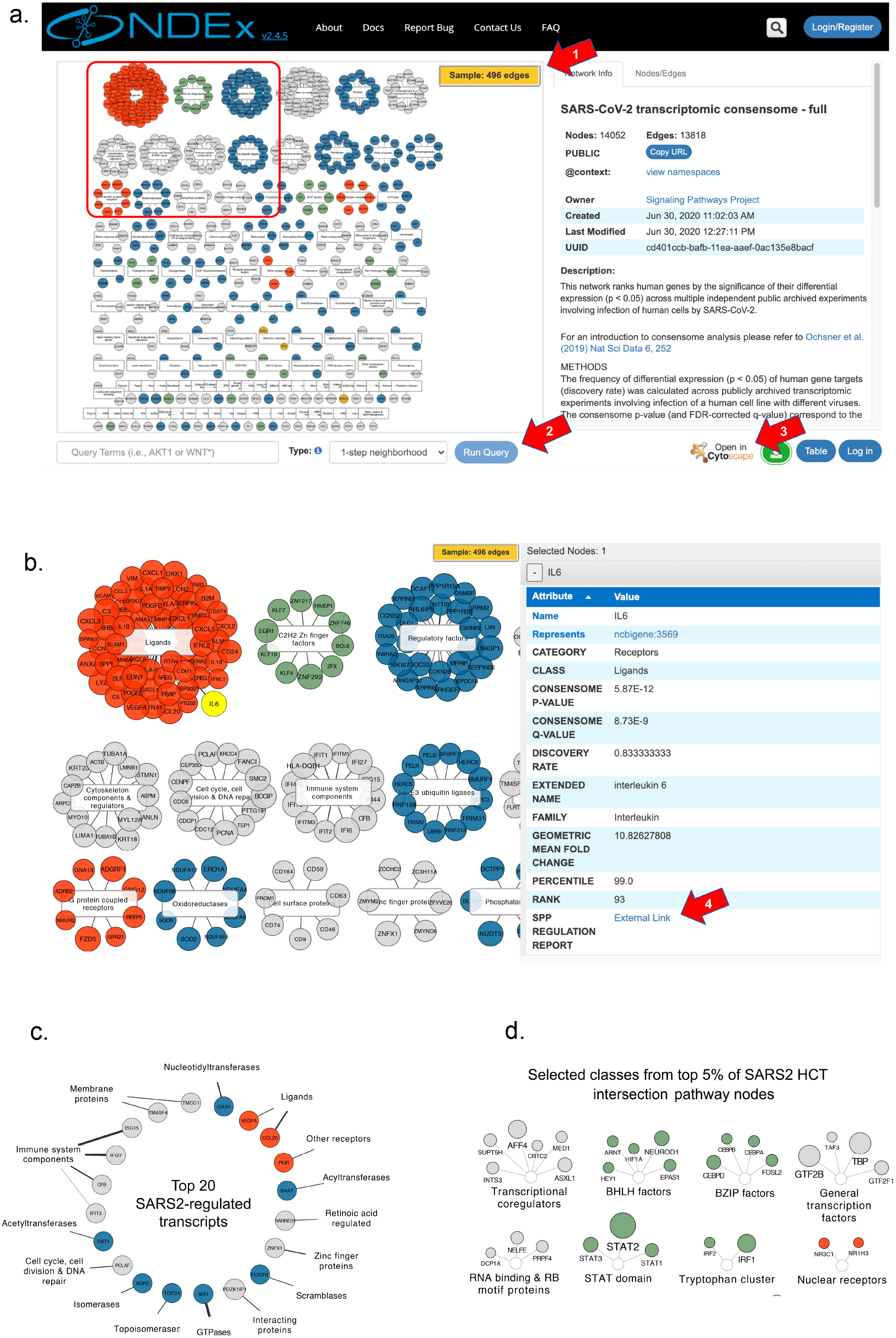
Visualization of viral consensomes and HCT intersection networks in the NDEx repository. In all panels, transcripts (consensome networks; panels a, b & c) and nodes (HCT intersection network; panel d) are color-coded according to their category as follows: receptors (orange), enzymes (blue), transcription factors (green), ion channels (mustard) and co-nodes (grey). Additional contextual information is available in the description of each network on the NDEx site. Red arrows are explained in the text. **a.** Sample view of SARS2 consensome showing top 5% of transcripts. White rectangles represent classes to which transcripts have been mapped in the SPP biocuration pipeline^10^. Orange rectangle refers to the view in panel b. **b.** Zoomed-in view of orange rectangle in panel A. IL6 transcript is highlighted to show the contextual information available in the side panel. **c.** Top 20 ranked transcripts in the SARS2 consensome. Edge widths are proportional to the GMFC. **d.** Selected classes represented in the top 5% of nodes in the SARS2 ChIP-Seq HCT intersection network. Node circle size is inversely proportional to the intersection q-value.

Consensome GMFCs are signless with respect to direction of regulation^10^. Researchers can therefore follow the SPP link in the side panel (Fig. 10b, red arrow 4) to view the individual underlying experimental data points on the SPP site (Fig. 9c shows the example for *IFI27*). A network of the top 20 ranked transcripts in the SARS2 consensome (Fig. 10c) includes genes with known (*OAS1, MX1*^114^) and previously uncharacterized (*PDZKIP1, SAT1, TM4SF4*) transcriptional responses to SARS2 infection. Finally, to afford insight into pathway nodes whose gain or loss of function contributes to SARS2 infection-induced signaling, Figure 10d shows the top 5% ranked nodes in the SARS2 node HCT ChIP-Seq intersection network (see figshare File F1, section 7; see also Figs. 2 & 3 and accompanying discussion above). In this, as with all HCT intersection networks, node size is proportional to the q-value, such that the larger the circle, the lower the q-value, and the higher the confidence that a particular node or node family is involved in the transcriptional response to viral infection.

The visual organization of the NDEx interface offers insights into the impact of CoV infection on human cell signaling that are not readily appreciated in the current SPP interface. For example, it is readily apparent from the NDEx SARS2 consensome network (Fig. 10c; Table 1) that the single largest class of SARS2 HCTs encodes immunomodulatory ligands (OR = 4.6, *p* = 3.8 e-24, hypergeometric test), many of which are members of the cytokine and chemokine superfamilies. In contrast, although still overabundant (OR = 1.58, p = 6.8e-4, hypergeometric test), inflammatory ligands comprise a considerably smaller proportion of the SARS1 HCTs (Table 1). These data represent evidence that SARS2 infection is relatively efficient in modulating a transcriptional inflammatory response in host cells. Consistent with this hypothesis, and while this manuscript was under review, a study appeared showing induction of interferon-stimulated genes in COVID-19 patients was more robust than in response to SARS1 infection^115^.

## Discussion

An effective research community response to the impact of CoV infection on human health demands systematic exploration of the transcriptional interface between CoV infection and human cell signaling systems. It also demands routine access to computational analysis of existing datasets that is unhindered either by paywalls or by lack of the informatics training required to manipulate archived datasets in their unprocessed state. Moreover, the substantial logistical obstacles to high containment laboratory certification emphasize the need for fullest possible access to, and re-usability of, existing CoV infection datasets to focus and refine hypotheses prior to carrying out *in vivo* CoV infection experiments. Meta-analysis of existing datasets represents a powerful approach to establishing consensus transcriptional signatures – consensomes – which identify those human genes whose expression is most consistently and reproducibly impacted by CoV infection. Moreover, integrating these consensus transcriptional signatures with existing consensomes for cellular signaling pathway nodes can illuminate transcriptional convergence between CoV infection and human cell signaling nodes.

To this end, we generated a set of CoV infection consensomes that rank human genes by the reproducibility of their differential expression (*p* < 0.05) in response to infection of human cells by CoVs. Using HCT intersection analysis, we then computed the CoV consensomes against high confidence transcriptional signatures for a broad range of cellular signaling pathway nodes, affording investigators with a broad range of signaling interests an entrez into the study of CoV infection of human cells. Although other enrichment based pathway analysis tools exist^116^, HCT intersection analysis differs from these by computing against only genes that have the closest predicted regulatory relationships with upstream pathway nodes (i.e. HCTs). The use cases described here represent illustrative examples of the types of analyses that users are empowered to carry out in the CoV infection knowledgebase.

Previous network analyses across independent viral infection transcriptomic datasets, although valuable, have been limited to stand-alone studies^117,118^. Here, to enable access to the CoV consensomes and their >3,000,000 underlying data points by the broadest possible audience, we have integrated them into the SPP knowledgebase and NDEx repository to create a unique, federated environment for generating hypotheses around the impact of CoV infection on human cell signaling. NDEx provides users with the familiar look and feel of Cytoscape to reduce barriers of accessibility and provides for intuitive click-and-drag data mining strategies. Incorporation of the CoV data points into SPP places them in the context of millions more existing SPP data points documenting transcriptional regulatory relationships between human pathway nodes and their genomic targets. In doing so, we provide users with evidence for signaling nodes whose gain or loss of function in response to CoV infection gives rise to these transcriptional patterns. The transcriptional impact of viral infection is known to be an amalgam of host antiviral responses and co-option by viruses of the host signaling machinery in furtherance of its life cycle. It is hoped that dissection of these two distinct modalities in the context of CoV infection will be facilitated by the availability of the CoV consensomes in the SPP and NDEx knowledgebases.

The CoV consensomes have a number of limitations. Primarily, since they are predicated specifically on transcriptional regulatory technologies, they will assign low rankings to transcripts that may not be transcriptionally responsive to CoV infection, but whose encoded proteins nevertheless play a role in the cellular response. For example, *MASP2*, which encodes an important node in the response to CoV infection has either a very low consensome ranking (SARS1, MERS and IAV), or is absent entirely (SARS2), indicating that it is transcriptionally unresponsive to viral infection and likely activated at the protein level in response to upstream signals. This and similar instances therefore represent “false negatives” in the context of the impact of CoV infection on human cells. Another limitation of the transcriptional focus of the datasets is the absence of information on specific protein interactions and post-translational modifications, either viral-human or human-human, that give rise to the observed transcriptional responses. Although these can be inferred to some extent, the availability of existing^32,68,109^ and future proteomic and kinomic datasets will facilitate modeling of the specific signal transduction events giving rise to the downstream transcriptional responses. Finally, although detailed metadata are readily available on the underlying data points, the consensomes do not directly reflect the impact of variables such as tissue context or duration of infection on differential gene expression. As additional suitable archived datasets become available, we will be better positioned to generate more specific consensomes of this nature.

The human CoV and IAV consensomes and their underlying datasets are intended as “living” resources in SPP and NDEx that will be updated and versioned with appropriate datasets as resources permit. This will be particularly important in the case of SARS2, given the expanded budget that worldwide funding agencies are likely to allocate to research into the impact of this virus on human health. Incorporation of future datasets will allow for clarification of observations that are intriguing, but whose significance is currently unclear, such as the intersection between the CoV HCTs and those of the telomerase catalytic subunit (figshare File F2), as well as the enrichment of EMT genes among those with elevated rankings in the SARS2 consensome (Fig. 7). Although they are currently available on the SPP website, distribution of the CoV consensome data points via the SPP RESTful API^10^ will be essential for the research community to fully capitalize on this work. For example, several co-morbidities of SARS2 infection, including renal and gastrointestinal disorders, are within the mission of the National Institute of Diabetes, Digestive and Kidney Diseases. In an ongoing collaboration with the NIDDK Information Network (DKNET)^120^, the SPP API will make the CoV consensome data points available in a hypothesis generation environment that will enable users to model intersections of CoV infection-modulated host signaling with their own research areas of interest. We welcome feedback and suggestions from the research community for the future development of the CoV infection consensomes and HCT node intersection networks.

## Methods

Consistent with emerging NIH mandates on rigor and reproducibility, we have used the Research Resource Identifier (RRID) standard^121^ to identify key research resources of relevance to our study.

### Dataset biocuration

Datasets from Gene Expression Omnibus (SCR_005012) and Array Express (SCR_002964) were biocurated as previously described, with the incorporation of an additional classification of peptide ligands^122^ to supplement the existing mappings derived from the International Union of Pharmacology Guide To Pharmacology (SCR_013077).

### Dataset processing and consensome analysis

#### Array data processing

To process microarray expression data, we utilized the log2 summarized and normalized array feature expression intensities provided by the investigator and housed in GEO. These data are available in the corresponding “Series Matrix Files(s)”. The full set of summarized and normalized sample expression values were extracted and processed in the statistical program R. To calculate differential gene expression for investigator-defined experimental contrasts, we used the linear modeling functions from the Bioconductor limma analysis package Initially, a linear model was fitted to a group-means parameterization design matrix defining each experimental variable. Subsequently, we fitted a contrast matrix that recapitulated the sample contrasts of interest, in this case viral infection vs mock infection, producing fold-change and significance values for each array feature present on the array. The current BioConductor array annotation library was used for annotation of array identifiers. P values obtained from limma analysis were not corrected for multiple comparisons.

#### RNA-Seq data processing

To process RNA-Seq expression data, we utilized the aligned, un-normalized, gene summarized read count data provided by the investigator and housed in GEO. These data are available in the corresponding “Supplementary file” section of the GEO record. The full set of raw aligned gene read count values were extracted and processed in the statistical program R using the limma^123^ and edgeR analysis^124^ packages. Read count values were initially filtered to remove genes with low read counts. Gene read count values were passed to downstream analysis if all replicate samples from at least one experimental condition had cpm > 1. Sequence library normalization factors were calculated to apply scale normalization to the raw aligned read counts using the TMM normalization method implemented in the edgeR package followed by the voom function^125^ to convert the gene read count values to log2-cpm. The log2-cpm values were initially fit to a group-means parameterization design matrix defining each experimental variable. This was subsequently fit to a contrast design matrix that recapitulates the sample contrasts of interest, in this case viral infection vs mock infection, producing fold-change and significance values for each aligned sequenced gene. If necessary, the current BioConductor human organism annotation library was used for annotation of investigator-provided gene identifiers. P values obtained from limma analysis were not corrected for multiple comparisons.

Differential expression values were committed to the consensome analysis pipeline as previously described^10^. Briefly, the consensome algorithm surveys each experiment across all datasets and ranks genes according to the frequency with which they are significantly differentially expressed. For each transcript, we counted the number of experiments where the significance for differential expression was ≤0.05, and then generated the binomial probability, referred to as the consensome p-value (CPV), of observing that many or more nominally significant experiments out of the number of experiments in which the transcript was assayed, given a true probability of 0.05. Genes were ranked firstly by CPV, then by geometric mean fold change (GMFC). A more detailed description of the transcriptomic consensome algorithm is available in a previous publication^10^. The consensomes and underlying datasets were loaded into an Oracle 13c database and made available on the SPP user interface as previously described^10^.

### Statistical analysis

High confidence transcript intersection analysis was performed using the Bioconductor GeneOverlap analysis package^17^ (SCR_018419) implemented in R. Briefly, given a whole set I of IDs and two sets A ∈ I and B ∈ I, and S = A ⋂ B, GeneOverlap calculates the significance of obtaining S. The problem is formulated as a hypergeometric distribution or contingency table, which is solved by Fisher’s exact test. *p*-values were adjusted for multiple testing by using the method of Benjamini & Hochberg to control the false discovery rate as implemented with the p.adjust function in R, to generate q-values. The universe for the intersection was set at a conservative estimate of the total number of transcribed (protein and non protein-coding) genes in the human genome (25,000)^126^, R code for analyzing the intersection between an investigator gene set and CoV consensome HCTs has been deposited in the SPP Github account. A two tailed two sample t-test assuming equal variance was used to compare the mean percentile ranking of the EMT (12 degrees of freedom) and E2F (14 degrees of freedom) signatures in the MERS, SARS1, SARS2 and IAV consensomes using the PRISM software package (SCR_005375).

### Consensome generation

The procedure for generating transcriptomic consensomes has been previously described^10^. To generate the ChIP-Seq consensomes, we first retrieved processed gene lists from ChIP-Atlas (SCR_015511), in which human genes are ranked based upon their average MACS2 occupancy across all publically archived datasets in which a given pathway node is the IP antigen. Of the three stringency levels available (10, 5 and 1 kb from the transcription start site), we selected the most stringent (1 kb). According to SPP convention^10^, we then mapped the IP antigen to its pathway node category, class and family, and the experimental cell line to its appropriate biosample physiological system and organ. We then organized the ranked lists into percentiles to generate the node ChIP-Seq consensomes. The 95^th^ percentiles of all consensomes (HCTs, high confidence transcriptional targets) was used as the input for the HCT intersection analysis.

### SPP web application

The SPP knowledgebase (SCR_018412) is a gene-centric Java Enterprise Edition 6, web-based application around which other gene, mRNA, protein and BSM data from external databases such as NCBI are collected. After undergoing semiautomated processing and biocuration as described above, the data and annotations are stored in SPP’s Oracle 13c database. RESTful web services exposing SPP data, which are served to responsively designed views in the user interface, were created using a Flat UI Toolkit with a combination of JavaScript, D3.JS, AJAX, HTML5, and CSS3. JavaServer Faces and PrimeFaces are the primary technologies behind the user interface. SPP has been optimized for Firefox 24+, Chrome 30+, Safari 5.1.9+, and Internet Explorer 9+, with validations performed in BrowserStack and load testing in LoadUIWeb. XML describing each dataset and experiment is generated and submitted to CrossRef (SCR_003217) to mint DOIs^10^.

## Supporting information

Figshare F1

Figshare F2

## Data availability

Important note on data availability: this paper refers to the first versions of the consensomes and HCT intersection networks based on the datasets available at the time of publication. As additional CoV infection datasets are archived over time, we will make updated versions of the consensomes and HCT intersection analyses accessible in future releases. The entire set of experimental metadata is available in figshare File F1, section 1. Consensome data points are in figshare File F1, sections 2-5.

### SPP

SPP MERS^137^, SARS1^141^, SARS2^145^ and IAV^149^ consensomes, their underlying data points and metadata, as well as original datasets, are freely accessible at https://ww.signalingpathways.org. Programmatic access to all underlying data points and their associated metadata are supported by a RESTful API at https://www.signalingpathways.org/docs/. All SPP datasets are biocurated versions of publically archived datasets, are formatted according to the recommendations of the FORCE11 Joint Declaration on Data Citation Principles, and are made available under a Creative Commons CC BY 4.0 license. The original datasets are available are linked to from the corresponding SPP datasets using the original repository accession identifiers. These identifiers are for transcriptomic datasets, the Gene Expression Omnibus (GEO) Series (GSE); and for cistromic/ChIP-Seq datasets, the NCBI Sequence Read Archive (SRA) study identifier (SRP). SPP consensomes are assigned DOIs as shown in Table 1.

### NDEx

NDEx versions of consensomes (MERS^138^, SARS1^142^, SARS2^146^ and IAV^150^) and node family (MERS^139^, SARS1^143^, SARS2^147^ and IAV^151^) and node (MERS^140^, SARS1^144^, SARS2^148^ and IAV^152^) HCT intersection networks are freely available in the NDEx repository and assigned DOIs as shown in Table 1. NDEx is a recommended repository for Scientific Data, Springer Nature and the PLOS family of journals and is registered on FAIRsharing.org; for additional info and documentation, please visit http://ndexbio.org. The official SPP account in NDEx is available at: https://bit.ly/30nN129.

## Code availability

SPP source code is available in the SPP GitHub account under a Creative Commons CC BY 4.0 license at https://github.com/signaling-pathways-project.

## Acknowledgements

This work was supported by the National Institute of Diabetes, Digestive and Kidney Diseases NIDDK Information Network (DK097748), the National Cancer Institute (CA125123, CA184427) and by the Brockman Medical Research Foundation. We acknowledge the assistance of Apollo McOwiti, Shijing Qu and Michael Dehart in making the datasets and consensomes available in the SPP knowledgebase. We thank all investigators who archive their datasets, without whom SPP would not be possible.

## Author contributions

**Dataset biocuration:** SO

**Data analysis:** SO, RP, NM

**Manuscript drafting:** NM

**Manuscript editing:** NM, RP, SO

## References

1. Takeda, K., Kaisho, T. & Akira, S. Toll-like receptors. Annu. Rev. Immunol. 21, 335–376 (2003).

2. Stark, G. R., Kerr, I. M., Williams, B. R., Silverman, R. H. & Schreiber, R. D. How cells respond to interferons. Annu. Rev. Biochem. 67, 227–64 (1998).

3. Darnell, J. E., Kerr, I. M. & Stark, G. R. Jak-STAT pathways and transcriptional activation in response to IFNs and other extracellular signaling proteins. Science (80-.). 264, 1415–1421 (1994).

4. DeDiego, M. L. et al. Severe acute respiratory syndrome coronavirus envelope protein regulates cell stress response and apoptosis. PLoS Pathog. 7, e1002315 (2011).

5. Josset, L. et al. Cell host response to infection with novel human coronavirus EMC predicts potential antivirals and important differences with SARS coronavirus. MBio 4, e00165–13 (2013).

6. Sims, A. C. et al. Release of severe acute respiratory syndrome coronavirus nuclear import block enhances host transcription in human lung cells. J. Virol. 87, 3885–3902 (2013).

7. Yoshikawa, T. et al. Dynamic innate immune responses of human bronchial epithelial cells to severe acute respiratory syndrome-associated coronavirus infection. PLoS One 5, e8729 (2010).

8. Blanco-Melo, D. et al. Imbalanced Host Response to SARS-CoV-2 Drives Development of COVID-19. Cell 181, 1036–1045.e9 (2020).

9. Lamers, M. M. et al. SARS-CoV-2 productively infects human gut enterocytes. Science https://doi.org/10.1126/science.abc1669 (2020).

10. Ochsner, S. A. et al. The Signaling Pathways Project, an integrated ‘omics knowledgebase for mammalian cellular signaling pathways. Sci. data 6, 252 (2019).

11. Aevermann, B. D. et al. A comprehensive collection of systems biology data characterizing the host response to viral infection. Sci. data 1, 140033 (2014).

12. Schneider, W. M., Chevillotte, M. D. & Rice, C. M. Interferon-stimulated genes: a complex web of host defenses. Annu. Rev. Immunol. 32, 513–545 (2014).

13. Doyle, T. et al. The interferon-inducible isoform of NCOA7 inhibits endosome-mediated viral entry. Nat. Microbiol. 3, 1369–1376 (2018).

14. Chapgier, A. et al. A partial form of recessive STAT1 deficiency in humans. J. Clin. Invest. 119, 1502–1514 (2009).

15. Gruter, P. et al. TAP, the human homolog of Mex67p, mediates CTE-dependent RNA export from the nucleus. Mol. Cell 1, 649–659 (1998).

16. Wei, J. et al. Genome-wide CRISPR screen reveals host genes that regulate SARS-CoV-2 infection. bioRxiv 2020.06.16.155101 https://doi.org/10.1101/2020.06.16.155101 (2020).

17. Shen, L. & Sinai, M. GeneOverlap: GeneOverlap: Test and visualize gene overlaps. R package version 1.24.0 http://shenlab-sinai.github.io/shenlab-sinai/ (2020).

18. Totura, A. L. et al. Toll-Like Receptor 3 Signaling via TRIF Contributes to a Protective Innate Immune Response to Severe Acute Respiratory Syndrome Coronavirus Infection. MBio 6, e00638–15 (2015).

19. Hensley, L. E. et al. Interferon-beta 1a and SARS coronavirus replication. Emerg. Infect. Dis. 10, 317–319 (2004).

20. Appelberg, S. et al. Dysregulation in mTOR/HIF-1 signaling identified by proteo-transcriptomics of SARS-CoV-2 infected cells. bioRxiv https://doi.org/10.1101/2020.04.30.070383 (2020).

21. Wang, W. et al. Up-regulation of IL-6 and TNF-alpha induced by SARS-coronavirus spike protein in murine macrophages via NF-kappaB pathway. Virus Res. 128, 1–8 (2007).

22. Zheng, K., Kitazato, K. & Wang, Y. Viruses exploit the function of epidermal growth factor receptor. Rev. Med. Virol. 24, 274–286 (2014).

23. Ng, S. S. M., Li, A., Pavlakis, G. N., Ozato, K. & Kino, T. Viral infection increases glucocorticoid-induced interleukin-10 production through ERK-mediated phosphorylation of the glucocorticoid receptor in dendritic cells: potential clinical implications. PLoS One 8, e63587 (2013).

24. Ostler, J. B., Harrison, K. S., Schroeder, K., Thunuguntla, P. & Jones, C. The Glucocorticoid Receptor (GR) Stimulates Herpes Simplex Virus 1 Productive Infection, in Part Because the Infected Cell Protein 0 (ICP0) Promoter Is Cooperatively Transactivated by the GR and Kruppel-Like Transcription Factor 15. J. Virol. 93, (2019).

25. Hayward, S. D. Viral interactions with the Notch pathway. Semin. Cancer Biol. 14, 387–396 (2004).

26. Ito, T. et al. The critical role of Notch ligand Delta-like 1 in the pathogenesis of influenza A virus (H1N1) infection. PLoS Pathog. 7, e1002341 (2011).

27. Rizzo, P. et al. COVID-19 in the heart and the lungs: could we ‘Notch’ the inflammatory storm? Basic research in cardiology vol. 115 31 (2020).

28. Wang, S. et al. Xenobiotic pregnane X receptor (PXR) regulates innate immunity via activation of NLRP3 inflammasome in vascular endothelial cells. J. Biol. Chem. 289, 30075–30081 (2014).

29. Marik, P. E., Kory, P. & Varon, J. Does vitamin D status impact mortality from SARS-CoV-2 infection? Medicine in drug discovery https://doi.org/10.1016/j.medidd.2020.100041 (2020).

30. Rhodes, J. M., Subramanian, S., Laird, E. & Kenny, R. A. Editorial: low population mortality from COVID-19 in countries south of latitude 35 degrees North supports vitamin D as a factor determining severity. Alimentary pharmacology & therapeutics https://doi.org/10.1111/apt.15777 (2020).

31. Ledford, H. Coronavirus breakthrough: dexamethasone is first drug shown to save lives. Nature https://doi.org/10.1038/d41586-020-01824-5 (2020).

32. Stukalov, A. et al. Multi-level proteomics reveals host-perturbation strategies of SARS-CoV-2 and SARS-CoV. bioRxiv 2020.06.17.156455 https://doi.org/10.1101/2020.06.17.156455 (2020).

33. Poppe, M. et al. The NF-kappaB-dependent and - independent transcriptome and chromatin landscapes of human coronavirus 229E-infected cells. PLoS Pathog. 13, e1006286 (2017).

34. Ludwig, S. & Planz, O. Influenza viruses and the NF-kappaB signaling pathway - towards a novel concept of antiviral therapy. Biol. Chem. 389, 1307–1312 (2008).

35. Ruckle, A. et al. The NS1 protein of influenza A virus blocks RIG-I-mediated activation of the noncanonical NF-kappaB pathway and p52/RelB-dependent gene expression in lung epithelial cells. J. Virol. 86, 10211–10217 (2012).

36. Chiang, H.-S. & Liu, H. M. The Molecular Basis of Viral Inhibition of IRF- and STAT-Dependent Immune Responses. Front. Immunol. 9, 3086 (2018).

37. Frieman, M. et al. Severe acute respiratory syndrome coronavirus ORF6 antagonizes STAT1 function by sequestering nuclear import factors on the rough endoplasmic reticulum/Golgi membrane. J. Virol. 81, 9812–9824 (2007).

38. Garcia-Sastre, A. et al. The role of interferon in influenza virus tissue tropism. J. Virol. 72, 8550–8558 (1998).

39. Blaszczyk, K. et al. The unique role of STAT2 in constitutive and IFN-induced transcription and antiviral responses. Cytokine Growth Factor Rev. 29, 71–81 (2016).

40. Boudewijns, R. et al. STAT2 signaling as double-edged sword restricting viral dissemination but driving severe pneumonia in SARS-CoV-2 infected hamsters. bioRxiv https://doi.org/10.1101/2020.04.23.056838 (2020).

41. Haas, D. A. et al. Viral targeting of TFIIB impairs de novo polymerase II recruitment and affects antiviral immunity. PLoS Pathog. 14, e1006980 (2018).

42. Gelev, V. et al. A new paradigm for transcription factor TFIIB functionality. Sci. Rep. 4, 3664 (2014).

43. Haviv, I., Shamay, M., Doitsh, G. & Shaul, Y. Hepatitis B virus pX targets TFIIB in transcription coactivation. Mol. Cell. Biol. 18, 1562–1569 (1998).

44. Bellail, A. C., Olson, J. J. & Hao, C. SUMO1 modification stabilizes CDK6 protein and drives the cell cycle and glioblastoma progression. Nat. Commun. 5, 4234 (2014).

45. Grossel, M. J. & Hinds, P. W. Beyond the cell cycle: a new role for Cdk6 in differentiation. J. Cell. Biochem. 97, 485–493 (2006).

46. Kaldis, P., Ojala, P. M., Tong, L., Makela, T. P. & Solomon, M. J. CAK-independent activation of CDK6 by a viral cyclin. Mol. Biol. Cell 12, 3987–3999 (2001).

47. Pauls, E. et al. Cell cycle control and HIV-1 susceptibility are linked by CDK6-dependent CDK2 phosphorylation of SAMHD1 in myeloid and lymphoid cells. J. Immunol. 193, 1988–1997 (2014).

48. Cingoz, O. & Goff, S. P. Cyclin-dependent kinase activity is required for type I interferon production. Proc. Natl. Acad. Sci. U. S. A. 115, E2950–E2959 (2018).

49. Hennessy, E. J., Sheedy, F. J., Santamaria, D., Barbacid, M. & O’Neill, L. A. J. Toll-like receptor-4 (TLR4) down-regulates microRNA-107, increasing macrophage adhesion via cyclin-dependent kinase 6. J. Biol. Chem. 286, 25531–25539 (2011).

50. Handschick, K. et al. Cyclin-dependent kinase 6 is a chromatin-bound cofactor for NF-kappaB-dependent gene expression. Mol. Cell 53, 193–208 (2014).

51. Zaborowska, J., Isa, N. F. & Murphy, S. P-TEFb goes viral. Bioessays 38 Suppl 1, S75–85 (2016).

52. Wobbe, C. R. et al. In vitro replication of DNA containing either the SV40 or the polyoma origin. Philos. Trans. R. Soc. Lond. B. Biol. Sci. 317, 439–453 (1987).

53. Takahashi, K. et al. DNA topoisomerase 1 facilitates the transcription and replication of the Ebola virus genome. J. Virol. 87, 8862–8869 (2013).

54. Li, W. et al. Angiotensin-converting enzyme 2 is a functional receptor for the SARS coronavirus. Nature 426, 450–454 (2003).

55. Hoffmann, M. et al. SARS-CoV-2 Cell Entry Depends on ACE2 and TMPRSS2 and Is Blocked by a Clinically Proven Protease Inhibitor. Cell 181, 271–280.e8 (2020).

56. Verdecchia, P., Cavallini, C., Spanevello, A. & Angeli, F. The pivotal link between ACE2 deficiency and SARS-CoV-2 infection. Eur. J. Intern. Med. https://doi.org/10.1016/j.ejim.2020.04.037 (2020).

57. Fadason, T. et al. A transcription regulatory network within the ACE2 locus may promote a pro-viral environment for SARS-CoV-2 by modulating expression of host factors. bioRxiv https://doi.org/10.1101/2020.04.14.042002 (2020).

58. Ziegler, C. G. K. et al. SARS-CoV-2 Receptor ACE2 Is an Interferon-Stimulated Gene in Human Airway Epithelial Cells and Is Detected in Specific Cell Subsets across Tissues. Cell https://doi.org/10.1016/j.cell.2020.04.035 (2020).

59. Zhu, H. et al. Clinical analysis of 10 neonates born to mothers with 2019-nCoV pneumonia. Transl. Pediatr. 9, 51–60 (2020).

60. Sappenfield, E., Jamieson, D. J. & Kourtis, A. P. Pregnancy and susceptibility to infectious diseases. Infect. Dis. Obstet. Gynecol. 2013, 752852 (2013).

61. Siston, A. M. et al. Pandemic 2009 influenza A(H1N1) virus illness among pregnant women in the United States. JAMA 303, 1517–1525 (2010).

62. Pastva, A., Estell, K., Schoeb, T. R. & Schwiebert, L. M. RU486 blocks the anti-inflammatory effects of exercise in a murine model of allergen-induced pulmonary inflammation. Brain. Behav. Immun. 19, 413–422 (2005).

63. Breslin, N. et al. COVID-19 infection among asymptomatic and symptomatic pregnant women: Two weeks of confirmed presentations to an affiliated pair of New York City hospitals. Am. J. Obstet. Gynecol. MFM 100118 https://doi.org/10.1016/j.ajogmf.2020.100118 (2020).

64. Sutton, D., Fuchs, K., D’Alton, M. & Goffman, D. Universal Screening for SARS-CoV-2 in Women Admitted for Delivery. The New England journal of medicine https://doi.org/10.1056/NEJMc2009316 (2020).

65. Bazer, F. W., Burghardt, R. C., Johnson, G. A., Spencer, T. E. & Wu, G. Interferons and progesterone for establishment and maintenance of pregnancy: interactions among novel cell signaling pathways. Reprod. Biol. 8, 179–211 (2008).

66. Richards, J. S., Russell, D. L., Ochsner, S. & Espey, L. L. Ovulation: New dimensions and new regulators of the inflammatory-like response. Annual Review of Physiology vol. 64 (2002).

67. Ghandehari, S. Progesterone for the Treatment of COVID-19 in Hospitalized Men. (2020).

68. Gordon, D. E. et al. A SARS-CoV-2 protein interaction map reveals targets for drug repurposing. Nature https://doi.org/10.1038/s41586-020-2286-9 (2020).

69. Lamouille, S., Xu, J. & Derynck, R. Molecular mechanisms of epithelial-mesenchymal transition. Nat. Rev. Mol. Cell Biol. 15, 178–196 (2014).

70. Hill, C., Jones, M. G., Davies, D. E. & Wang, Y. Epithelial-mesenchymal transition contributes to pulmonary fibrosis via aberrant epithelial/fibroblastic cross-talk. J. lung Heal. Dis. 3, 31–35 (2019).

71. Li, H. et al. Alveolar epithelial cells undergo epithelial-mesenchymal transition in acute interstitial pneumonia: a case report. BMC Pulm. Med. 14, 67 (2014).

72. Gouda, M. M., Shaikh, S. B. & Bhandary, Y. P. Inflammatory and Fibrinolytic System in Acute Respiratory Distress Syndrome. Lung 196, 609–616 (2018).

73. George, P., Wells, A. & Jenkins, G. Pulmonary fibrosis and COVID-19: the potential role for antifibrotic therapy. Lancet https://doi.org/10.1016/S2213-2600(20)30225-3 (2020).

74. Adair, L. B. 2nd & Ledermann, E. J. Chest CT Findings of Early and Progressive Phase COVID-19 Infection from a US Patient. Radiology case reports https://doi.org/10.1016/j.radcr.2020.04.031 (2020).

75. Zhou, P. et al. A pneumonia outbreak associated with a new coronavirus of probable bat origin. Nature 579, 270–273 (2020).

76. Bolós, V. et al. The transcription factor Slug represses E-cadherin expression and induces epithelial to mesenchymal transitions: a comparison with Snail and E47 repressors. J. Cell Sci. 116, 499–511 (2003).

77. Yang, J. et al. HIF-2α promotes epithelial-mesenchymal transition through regulating Twist2 binding to the promoter of E-cadherin in pancreatic cancer. J. Exp. Clin. Cancer Res. 35, 26 (2016).

78. Kobayashi, W. & Ozawa, M. The transcription factor LEF-1 induces an epithelial-mesenchymal transition in MDCK cells independent of β-catenin. Biochem. Biophys. Res. Commun. 442, 133–138 (2013).

79. Xu, J., Lamouille, S. & Derynck, R. TGF-beta-induced epithelial to mesenchymal transition. Cell Res. 19, 156–172 (2009).

80. Zhao, M., Kong, L., Liu, Y. & Qu, H. dbEMT: an epithelial-mesenchymal transition associated gene resource. Sci. Rep. 5, 11459 (2015).

81. Moheimani, F. et al. Influenza A virus infection dysregulates the expression of microRNA-22 and its targets; CD147 and HDAC4, in epithelium of asthmatics. Respir. Res. 19, 145 (2018).

82. Xia, L., Dai, L., Yu, Q. & Yang, Q. Persistent Transmissible Gastroenteritis Virus Infection Enhances Enterotoxigenic Escherichia coli K88 Adhesion by Promoting Epithelial-Mesenchymal Transition in Intestinal Epithelial Cells. J. Virol. 91, (2017).

83. Yang, J.-X. et al. Lipoxin A(4) ameliorates lipopolysaccharide-induced lung injury through stimulating epithelial proliferation, reducing epithelial cell apoptosis and inhibits epithelial-mesenchymal transition. Respir. Res. 20, 192 (2019).

84. Diaz-Pina, G. et al. The Role of ADAR1 and ADAR2 in the Regulation of miRNA-21 in Idiopathic Pulmonary Fibrosis. Lung 196, 393–400 (2018).

85. Vukmirovic, M. et al. Identification and validation of differentially expressed transcripts by RNA-sequencing of formalin-fixed, paraffin-embedded (FFPE) lung tissue from patients with Idiopathic Pulmonary Fibrosis. BMC Pulm. Med. 17, 15 (2017).

86. Gao, F., Kinnula, V. L., Myllarniemi, M. & Oury, T. D. Extracellular superoxide dismutase in pulmonary fibrosis. Antioxid. Redox Signal. 10, 343–354 (2008).

87. Kato, N. et al. Basigin/CD147 promotes renal fibrosis after unilateral ureteral obstruction. Am. J. Pathol. 178, 572–579 (2011).

88. Ruster, C. & Wolf, G. Angiotensin II as a morphogenic cytokine stimulating renal fibrogenesis. J. Am. Soc. Nephrol. 22, 1189–1199 (2011).

89. Wang, C. et al. CD147 Induces Epithelial-to-Mesenchymal Transition by Disassembling Cellular Apoptosis Susceptibility Protein/E-Cadherin/beta-Catenin Complex in Human Endometriosis. Am. J. Pathol. 188, 1597–1607 (2018).

90. Stewart, C. A. et al. SARS-CoV-2 infection induces EMT-like molecular changes, including ZEB1-mediated repression of the viral receptor ACE2, in lung cancer models. bioRxiv 2020.05.28.122291 https://doi.org/10.1101/2020.05.28.122291 (2020).

91. Shan, B., Farmer, A. A. & Lee, W. H. The molecular basis of E2F-1/DP-1-induced S-phase entry and apoptosis. Cell growth Differ. Mol. Biol. J. Am. Assoc. Cancer Res. 7, 689–697 (1996).

92. Harbour, J. W. & Dean, D. C. The Rb/E2F pathway: expanding roles and emerging paradigms. Genes Dev. 14, 2393–2409 (2000).

93. Reimer, D. et al. E2F3a is critically involved in epidermal growth factor receptor-directed proliferation in ovarian cancer. Cancer Res. 70, 4613–4623 (2010).

94. Classon, M. & Harlow, E. The retinoblastoma tumour suppressor in development and cancer. Nat. Rev. Cancer 2, 910–917 (2002).

95. Diril, M. K. et al. Cyclin-dependent kinase 1 (Cdk1) is essential for cell division and suppression of DNA re-replication but not for liver regeneration. Proc. Natl. Acad. Sci. U. S. A. 109, 3826–3831 (2012).

96. Shivji, K. K., Kenny, M. K. & Wood, R. D. Proliferating cell nuclear antigen is required for DNA excision repair. Cell 69, 367–374 (1992).

97. Borlado, L. R. & Méndez, J. CDC6: from DNA replication to cell cycle checkpoints and oncogenesis. Carcinogenesis 29, 237–243 (2008).

98. Holt, S. V et al. Silencing Cenp-F weakens centromeric cohesion, prevents chromosome alignment and activates the spindle checkpoint. J. Cell Sci. 118, 4889–4900 (2005).

99. Vanden Bosch, A. et al. NuSAP is essential for chromatin-induced spindle formation during early embryogenesis. J. Cell Sci. 123, 3244–3255 (2010).

100. Engeland, K. Cell cycle arrest through indirect transcriptional repression by p53: I have a DREAM. Cell Death Differ. 25, 114–132 (2018).

101. Wang, N. et al. Increased expression of RRM2 by human papillomavirus E7 oncoprotein promotes angiogenesis in cervical cancer. Br. J. Cancer 110, 1034–1044 (2014).

102. Li, Y.-Y., Wang, L. & Lu, C.-D. An E2F site in the 5’-promoter region contributes to serum-dependent up-regulation of the human proliferating cell nuclear antigen gene. FEBS Lett. 544, 112–118 (2003).

103. Furukawa, Y., Terui, Y., Sakoe, K., Ohta, M. & Saito, M. The role of cellular transcription factor E2F in the regulation of cdc2 mRNA expression and cell cycle control of human hematopoietic cells. J. Biol. Chem. 269, 26249–26258 (1994).

104. Yuan, X. et al. SARS coronavirus 7a protein blocks cell cycle progression at G0/G1 phase via the cyclin D3/pRb pathway. Virology 346, 74–85 (2006).

105. Fan, Y., Sanyal, S. & Bruzzone, R. Breaking Bad: How Viruses Subvert the Cell Cycle. Front. Cell. Infect. Microbiol. 8, 396 (2018).

106. Gasser, S. & Raulet, D. H. The DNA damage response arouses the immune system. Cancer Res. 66, 3959–3962 (2006).

107. Kong, L.-J., Chang, J. T., Bild, A. H. & Nevins, J. R. Compensation and specificity of function within the E2F family. Oncogene 26, 321–327 (2007).

108. To, B. & Andrechek, E. R. Transcription factor compensation during mammary gland development in E2F knockout mice. PLoS One 13, e0194937 (2018).

109. M. Bouhaddou & Krogan, N. The Global Phosphorylation Landscape of SARS-CoV-2 Infection. Cell https://doi.org/10.1016/j.cell.2020.06.034 (2020).

110. Pillich, R. T., Chen, J., Rynkov, V., Welker, D. & Pratt, D. NDEx: A Community Resource for Sharing and Publishing of Biological Networks. Methods Mol. Biol. 1558, 271–301 (2017).

111. Pratt, D. et al. NDEx 2.0: A Clearinghouse for Research on Cancer Pathways. Cancer Res. 77, e58–e61 (2017).

112. Shannon, P. et al. Cytoscape: a software environment for integrated models of biomolecular interaction networks. Genome Res. 13, 2498–2504 (2003).

113. Chen, X. et al. Detectable serum SARS-CoV-2 viral load (RNAaemia) is closely correlated with drastically elevated interleukin 6 (IL-6) level in critically ill COVID-19 patients. Clin. Infect. Dis. https://doi.org/10.1093/cid/ciaa449 (2020).

114. Sungnak, W. et al. SARS-CoV-2 entry factors are highly expressed in nasal epithelial cells together with innate immune genes. Nat. Med. 26, 681–687 (2020).

115. Zhou, Z. et al. Heightened Innate Immune Responses in the Respiratory Tract of COVID-19 Patients. Cell Host Microbe 27, 883–890.e2 (2020).

116. Cirillo, E., Parnell, L. D. & Evelo, C. T. A Review of Pathway-Based Analysis Tools That Visualize Genetic Variants. Front. Genet. 8, 174 (2017).

117. Mitchell, H. D. et al. A network integration approach to predict conserved regulators related to pathogenicity of influenza and SARS-CoV respiratory viruses. PLoS One 8, e69374 (2013).

118. Ackerman, E. E., Alcorn, J. F., Hase, T. & Shoemaker, J. E. A dual controllability analysis of influenza virus-host protein-protein interaction networks for antiviral drug target discovery. BMC Bioinformatics 20, 297 (2019).

119. Wilk, C. M. Coronaviruses hijack the complement system. Nat. Rev. Immunol. https://doi.org/10.1038/s41577-020-0314-5 (2020).

120. Whetzel, P. L., Grethe, J. S., Banks, D. E. & Martone, M. E. The NIDDK Information Network: A Community Portal for Finding Data, Materials, and Tools for Researchers Studying Diabetes, Digestive, and Kidney Diseases. PLoS One 10, e0136206 (2015).

121. Bandrowski, A. E. & Martone, M. E. RRIDs: A Simple Step toward Improving Reproducibility through Rigor and Transparency of Experimental Methods. Neuron 90, 434–436 (2016).

122. Ramilowski, J. A. et al. A draft network of ligand-receptor-mediated multicellular signalling in human. Nat. Commun. 6, 7866 (2015).

123. Ritchie, M. E. et al. limma powers differential expression analyses for RNA-sequencing and microarray studies. Nucleic Acids Res. 43, e47 (2015).

124. Robinson, M. D., McCarthy, D. J. & Smyth, G. K. edgeR: a Bioconductor package for differential expression analysis of digital gene expression data. Bioinformatics 26, 139–140 (2010).

125. Law, C. W., Chen, Y., Shi, W. & Smyth, G. K. voom: Precision weights unlock linear model analysis tools for RNA-seq read counts. Genome Biol. 15, R29 (2014).

126. Pertea, M. et al. CHESS: a new human gene catalog curated from thousands of large-scale RNA sequencing experiments reveals extensive transcriptional noise. Genome Biol. 19, 208 (2018).

127. Oki, S. et al. ChIP-Atlas: a data-mining suite powered by full integration of public ChIP-seq data. EMBO Rep. 19, (2018).

128. Muzikar, K., Nickols, N. & Dervan, P. Analysis of the dexamethasone (Dex)-dependent transcriptome in A549 lung adenocarcinoma cells. Signaling Pathways Project Datasets. https://doi.org/10.1621/xigKzGn1se (2015).

129. Schulze-Gahmen, U. et al. The AFF4 scaffold binds human P-TEFb adjacent to HIV Tat. Elife 2, e00327 (2013).

130. Chalabi Hagkarim, N. et al. Degradation of a Novel DNA Damage Response Protein, Tankyrase 1 Binding Protein 1, following Adenovirus Infection. J. Virol. 92, (2018).

131. Yang, J. et al. Telomerase activation by Epstein-Barr virus latent membrane protein 1 is associated with c-Myc expression in human nasopharyngeal epithelial cells. J. Exp. Clin. Cancer Res. 23, 495–506 (2004).

132. Klingelhutz, A. J., Foster, S. A. & McDougall, J. K. Telomerase activation by the E6 gene product of human papillomavirus type 16. Nature 380, 79–82 (1996).

133. Gewin, L., Myers, H., Kiyono, T. & Galloway, D. A. Identification of a novel telomerase repressor that interacts with the human papillomavirus type-16 E6/E6-AP complex. Genes Dev. 18, 2269–2282 (2004).

134. Yin, L., Hubbard, A. K. & Giardina, C. NF-kappa B regulates transcription of the mouse telomerase catalytic subunit. J. Biol. Chem. 275, 36671–36675 (2000).

135. Akiyama, M. et al. Nuclear factor-kappaB p65 mediates tumor necrosis factor alpha-induced nuclear translocation of telomerase reverse transcriptase protein. Cancer Res. 63, 18–21 (2003).

136. Ghosh, A. et al. Telomerase directly regulates NF-kappaB-dependent transcription. Nat. Cell Biol. 14, 1270–1281 (2012).

137. Signaling Pathways Project Datasets. The MERS-CoV transcriptomic consensome. https://doi.org/10.1621/jgxM527b8s.1 (2020)

138. The Network Data Exchange. The MERS-CoV transcriptomic consensome network. https://doi.org/10.18119/N9QG7S (2020)

139. The Network Data Exchange. MERS-CoV node family high confidence transcriptional target intersection analysis network. https://doi.org/10.18119/N9PG63 (2020)

140. The Network Data Exchange. MERS-CoV node high confidence transcriptional target intersection analysis network. https://doi.org/10.18119/N96G6R (2020)

141. Signaling Pathways Project Datasets. The SARS-CoV-1 transcriptomic consensome. https://doi.org/10.1621/jgxM527b8s.1 (2020)

142. The Network Data Exchange. The SARS-CoV-1 transcriptomic consensome network. https://doi.org/10.18119/N9QG7S (2020)

143. The Network Data Exchange. SARS-CoV-1 node family high confidence transcriptional target intersection analysis network. https://doi.org/10.18119/N9PG63 (2020)

144. The Network Data Exchange. SARS-CoV-1 node high confidence transcriptional target intersection analysis network. https://doi.org/10.18119/N96G6R (2020)

145. Signaling Pathways Project Datasets. The SARS-CoV-2 transcriptomic consensome. https://doi.org/10.1621/jgxM527b8s.1 (2020)

146. The Network Data Exchange. The SARS-CoV-2 transcriptomic consensome network. https://doi.org/10.18119/N9QG7S (2020)

147. The Network Data Exchange. SARS-CoV-2 node family high confidence transcriptional target intersection analysis network. https://doi.org/10.18119/N9PG63 (2020)

148. The Network Data Exchange. SARS-CoV-2 node high confidence transcriptional target intersection analysis network. https://doi.org/10.18119/N96G6R (2020)

149. Signaling Pathways Project Datasets. The IAV transcriptomic consensome. https://doi.org/10.1621/jgxM527b8s.1 (2020)

150. The Network Data Exchange. The IAV transcriptomic consensome network. https://doi.org/10.18119/N9QG7S (2020)

151. The Network Data Exchange. IAV node family high confidence transcriptional target intersection analysis network. https://doi.org/10.18119/N9PG63 (2020)

152. The Network Data Exchange. IAV node high confidence transcriptional target intersection analysis network. https://doi.org/10.18119/N96G6R (2020)

