## Supplementary material for "Consensus transcriptional regulatory networks of coronavirus-infected human cells": Figshare F2

### Section 1. Co-node high confidence transcript (HCT) overlap analysis

Please refer to Supplementary file 1, section 7. We have coined the term “co-nodes” as a catch-all for cellular factors that are not members of the three principal signaling pathway node categories (receptors, enzymes, ion channels and transcription factors[1]. The breadth of functions encompassed by members of this category reflects the diverse mechanisms employed both by viruses to infect and propagate in cells, as well as by hosts to mount an efficient immune response. Consistent with its characterized role in the recruitment of p-TEFB by IV-1 Tat protein [2] for example, we observed consistently strong enrichment of the AFF4 CC95 in all viral TC95s. The targeting of CNOT3 for degradation in response to adenoviral infection [3] is reflected in the significant overlap between its HCTs and the viral HCTs, particularly in the case of SARS2.

### Section 2. Supplementary hypothesis generation use case: evidence for a role for the telomerase catalytic subunit TERT in the interferon response to CoV and IAV infection

Although telomerase activation has been well characterized in the context of cell immortalization by human tumor virus infection [4–6], no connection has previously been made between CoV or IAV infection and telomerase. We were therefore intrigued to observe robust overlap between all viral HCTs and those of the telomerase catalytic subunit TERT (SARS1, q = 7e-22; SARS2, q = 2e-12; MERS, 3.3e-16; IAV, 4e-28; Supplementary file 1, section 7). In support of this finding, NFkB signaling has been shown to induce expression [7] and nuclear translocation [8] of TERT, and direct co-regulation by telomerase of NFkB-dependent transcription has been linked to chronic inflammation [9]. Inspecting the consensome underlying data points (data not shown) we found that the *TERT* gene was not transcriptionally induced in response to infection by any of the CoVs, indicating that the overlap between its HCTs and those of the CoVs might occur in response to an upstream regulatory signal. If functional interactions between TERT and inflammatory nodes did indeed take place in response to CoV infection, we anticipated that this would be reflected in close agreement regarding the direction of differential expression of CoV infection-regulated genes in response to perturbation of TERT on the one hand, and on the other, to stimulation of the classic viral response IFNRs. To test this hypothesis, we took the same set of 20 ISGs referred to previously (Section 1) and compared their direction of regulation across all experiments encompassing the CoV, TERT and IFNR consensomes. For reference, the TERT consensome is provided in Supplementary file, section 12. With respect to the IFNR and TERT data points, we observed a nearly universal alignment in the direction of regulation of all genes tested with those in the CoV infection experiments (Supplementary Fig. 1), with agreement in the direction of regulation across 99% of the underlying probesets. We should note that of the 1859 $p < 0.05$ CoV infection ISG data points, we observed repression, rather than induction, in response to CoV infection in 303 (~15%) , which may be attributable to the impact of differences in cell type, cell cycle stage or virus incubation time across the independent experiments. Our results suggest the hypothesis that activation of telomerase is a component of the human response to CoV infection.

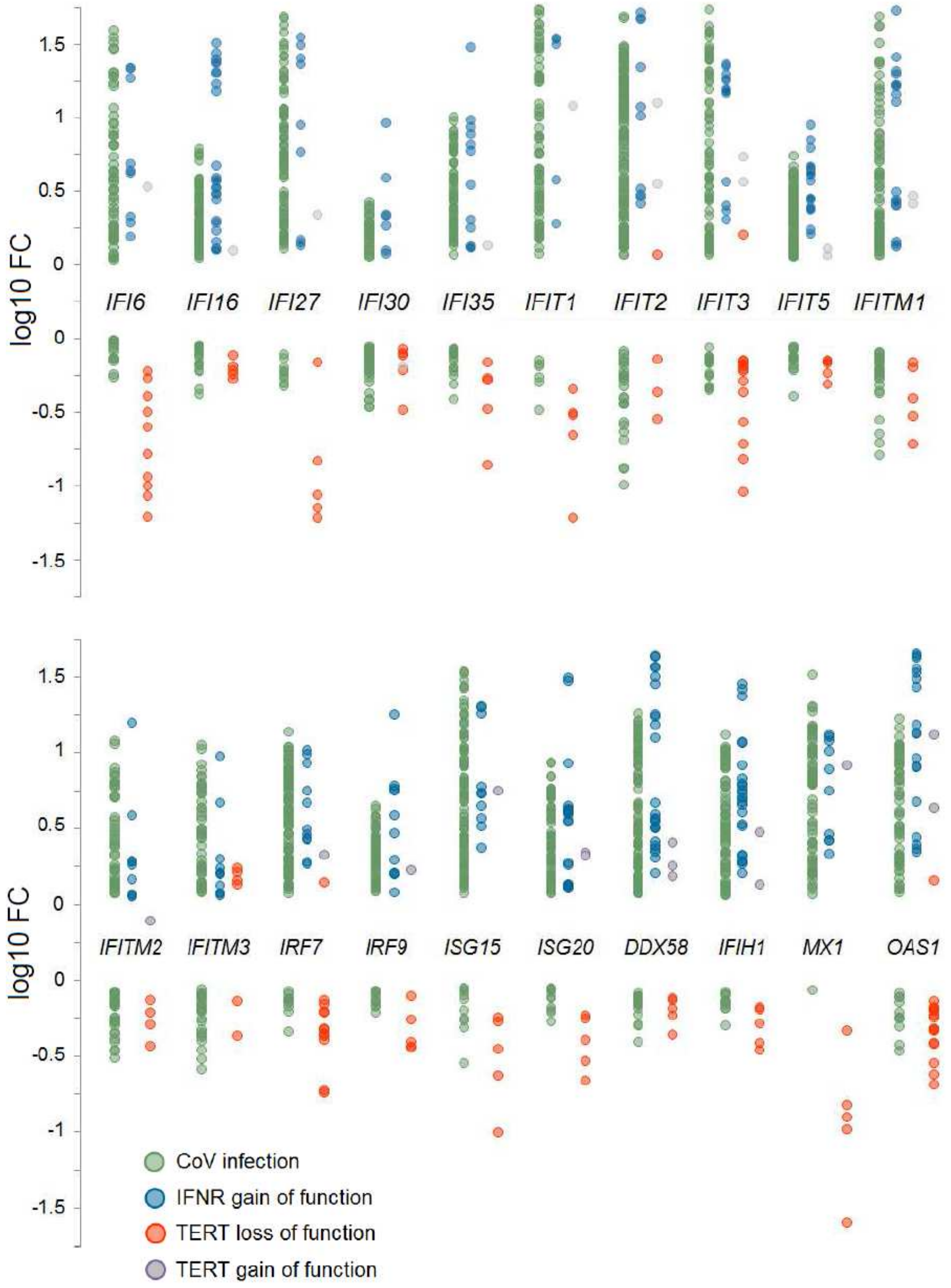


**Supplementary Figure 1. Conservation of polarity of differential expression in CoV, IFNR and TERT perturbation transcriptomic experiments across diverse canonical interferon-inducible viral response genes.** Positive/negative regulatory relationship of TERT with a transcriptional target was inferred from the design of the underlying experiments: this was loss of function for experiments in GSE77014 (MST132 inhibitor) and GSE60175 (shRNA), and gain of function in E-MEXP-563 (overexpression). All IFNR experiments were gain of function using characterized physiological ligands for members of the family. Refer to Supplementary File 1, section 1 for full information on the underlying experiments

1. Ochsner, S. A. *et al.* The Signaling Pathways Project, an integrated ’omics knowledgebase for mammalian  cellular signaling pathways. *Sci. data* **6**, 252 (2019).

2. Schulze-Gahmen, U. *et al.* The AFF4 scaffold binds human P-TEFb adjacent to HIV Tat. *Elife* **2**, e00327 (2013).

3. Chalabi Hagkarim, N. *et al.* Degradation of a Novel DNA Damage Response Protein, Tankyrase 1 Binding Protein 1, following Adenovirus Infection. *J. Virol.* **92**, (2018).

4. Yang, J. *et al.* Telomerase activation by Epstein-Barr virus latent membrane protein 1 is associated with c-Myc expression in human nasopharyngeal epithelial cells. *J. Exp. Clin. Cancer Res.* **23**, 495–506 (2004).

5. Klingelhutz, A. J., Foster, S. A. & McDougall, J. K. Telomerase activation by the E6 gene product of human papillomavirus type 16. *Nature* **380**, 79–82 (1996).

6. Gewin, L., Myers, H., Kiyono, T. & Galloway, D. A. Identification of a novel telomerase repressor that interacts with the human papillomavirus type-16 E6/E6-AP complex. *Genes Dev.* **18**, 2269–2282 (2004).

7. Yin, L., Hubbard, A. K. & Giardina, C. NF-kappa B regulates transcription of the mouse telomerase catalytic subunit. *J. Biol. Chem.* **275**, 36671–36675 (2000).

8. Akiyama, M. *et al.* Nuclear factor-kappaB p65 mediates tumor necrosis factor alpha-induced nuclear translocation of telomerase reverse transcriptase protein. *Cancer Res.* **63**, 18–21 (2003).

9. Ghosh, A. *et al.* Telomerase directly regulates NF-kappaB-dependent

transcription. *Nat. Cell Biol.* **14**, 1270–1281 (2012).
